# Viral Infections Drive Functional Remodeling of Ribosome-Associated Proteins

**DOI:** 10.64898/2026.01.29.701737

**Authors:** Thibault J.M. Sohier, Christelle Morris, David Cluet, Honglin Chen, Vincenzo Ruscica, Laura Guiguettaz, Anne-Marie Hesse, Sylvie Kieffer-Jaquinod, Shabaz Mohammed, Yohann Couté, Alfredo Castello, Emiliano P. Ricci

## Abstract

Viruses hijack host ribosomes to synthesize their proteins and have evolved various strategies to manipulate translation, including the association of numerous non-ribosomal proteins, known as ribosome-associated proteins (RAPs), with the core ribosome. However, current methods of capturing RAPs that associate transiently and/or in small amounts with ribosomes are suboptimal. To address this issue, we developed an enhanced ribosome affinity purification (eRAP) method that enables the capture of over 500 previously unreported RAPs. Using eRAP, we investigated whether the composition of ribosomes changes during infection by two very different viruses: herpes simplex virus 1 (HSV-1) and Sindbis virus (SINV). Our results revealed that viral infection does not substantially alter the composition of core ribosomes but does drive extensive remodeling of RAPs in a virus-specific manner. Functional analyses with SINV showed that several of these RAPs are critical for viral propagation. Additionally, we discovered that the RAP ASCC3 plays a novel role in promoting the expression of secretory proteins, including viral glycoproteins. This occurs independently of ASCC3’s canonical function of rescuing ribosome collisions. Together, our findings establish the ribosome as a dynamic regulatory hub that undergoes remodeling to achieve virus-specific translational output, highlighting the functional significance of ribosomal heterogeneity.

## Introduction

Ribosomes are sophisticated macromolecular complexes that translate messenger RNAs (mRNAs) into proteins. Eukaryotic ribosomes consist of four ribosomal RNAs (rRNAs) and approximately 80 ribosomal proteins (RPs), which are arranged into 60S and 40S subunits. Though long considered uniform “molecular machines”, ribosomes are now recognized as being more dynamic and heterogeneous than previously thought. Variability can arise at multiple levels, including rRNA modifications, RP stoichiometry, incorporation of RP paralogs, and post-translational modifications^1–4^. This heterogeneity is further shaped by the association of non-ribosomal proteins known as ribosome-associated proteins (RAPs), which provide an additional regulatory layer^5–7^. This diversity raises the possibility that ribosomes may execute specialized translational programs. However, the functional relevance of this heterogeneity remains debated^8^.

Viruses depend entirely on host ribosomes to synthesize their proteins, and are thus excellent model systems to explore ribosome remodeling. Viruses have indeed developed numerous strategies to subvert host translation, ranging from suppression of cap-dependent initiation to degradation of host mRNAs or selective engagement of RPs^9–12^. Viruses can also induce ribosome heterogeneity through post-translational modifications of RPs that favor viral translation^13–15^. Although some examples of virus-ribosome interactions have been documented^16^, it is unclear whether viral infection extensively remodel the ribosome-associated proteome to promote specific modes of translation. Answering this question could provide valuable insight into how ribosome heterogeneity is harnessed by physiological and pathological cues.

Here, we used two evolutionarily distant viruses: herpes simplex virus 1 (HSV-1), which is a nuclear double-stranded DNA virus; and Sindbis virus (SINV), which is a cytoplasmic positive-strand RNA alphavirus. By exploiting these two distinct infection strategies and genetic material, we aimed to reveal the breadth of ribosome reprogramming during infection. To profile dynamic changes in ribosome composition with high precision and depth, we developed enhanced ribosome affinity purification (eRAP), which enables the highly specific capture of cytoplasmic ribosomes with their associated proteins. eRAP unveiled that viral infection does not substantially alter the composition of core RPs but does drive extensive remodeling of RAPs in a virus-specific manner. Functional analyses revealed that several of these RAPs are critical for viral propagation. One such RAP is ASCC3, which unexpectedly promotes the expression of viral and cellular secretory proteins, independently of its canonical role in the ZNF598-mediated ribosome collision pathway. These findings establish viral infection as a powerful model for discovering new regulatory layers of gene expression by controlling ribosome composition, probing the functional significance of ribosome heterogeneity. Moreover, our study highlights the ribosome as a critical hub for host-virus interactions.

## Results

### eRAP, a powerful method to capture previously unreported ribosome-associated proteins

To study the effect of viral infection on ribosome interactome remodeling, we first developed eRAP. This method is designed to efficiently and specifically purify cytoplasmic ribosomes and their associated proteins, while excluding nuclear precursors and mitochondrial ribosomes. eRAP consists of endogenous tagging of an RP, mild formaldehyde crosslinking, lysis in presence of micrococcal nuclease (MNase), and quantitative proteomic analysis of the ribosome interactome in human cells (Figure 1A). The tagged RP was selected following two criteria: 1) accessibility of its N- or C-terminal ends on the ribosome surface, and 2) a low number of pseudogenes to ensure the specificity of CRISPR/Cas9-induced DNA breaks and recombination. We chose the RPS5/uS7 40S component as it fulfills these two critical requisites^17^ (Figure 1B). Furthermore, uS7 is present in stoichiometric amounts within both monosomes and polysomes, and it does not exhibit differential ribosome association in different mouse tissues^5,18^. The sequence encoding an 8xHis-Flag tag was inserted downstream of the methionine initiator codon of the *RPS5* gene in the human HEK293T cell line using CRISPR/Cas9-mediated genome editing (Figure 1C). A homozygous clone was derived to confirm that the 8xHis-Flag tag does not affect cell viability while ensuring homogeneous ribosome populations. Correct incorporation of 8xHis-Flag-tagged uS7 protein (HF-uS7) into the ribosome was verified by immunoblotting after separation of ribosome populations on sucrose gradients (Figure 1D). HF-uS7 was distributed similarly to uS3, another small ribosomal subunit protein, suggesting that the 8xHis-Flag tag did not interfere with ribosome assembly. Importantly, HF-uS7 was not detected in free fractions, ruling out extra-ribosomal functions.

**Figure 1.**
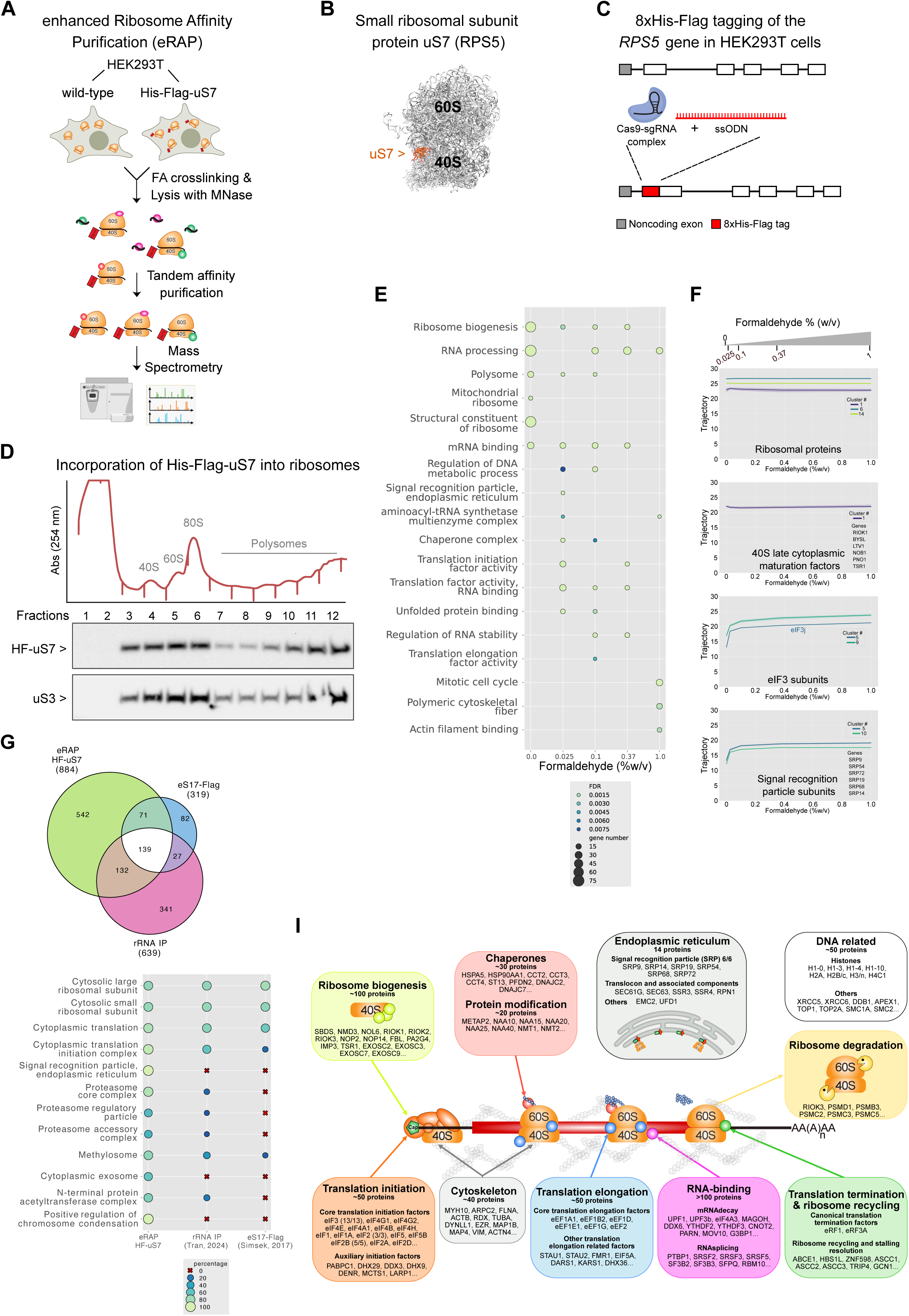
(A) Schematic of the enhanced ribosome affinity purification (eRAP) method. It includes mild formaldehyde crosslinking, partial mRNA digestion using MNase, and tandem affinity purification of ribosomes from cells containing an endogenous double-tagged ribosomal protein. (B) The location of the uS7 ribosomal protein (RPS5) is shown in a structural model of the human 80S ribosome^17^ (PDB: 4UG0). uS7, which is colored orange, is positioned at the head of the 40S ribosomal subunit. (C) Schematic representation of tagging the endogenous uS7 ribosomal protein with an 8xHis-Flag tag in HEK293T cells. The CRISPR-Cas9 endonuclease system was used to insert the 8xHis-Flag tag sequence by homology-directed repair downstream of the RPS5 translation start codon. This process employed a single guide RNA (sgRNA) targeting the RPS5 locus and a single-stranded DNA donor template (ssODN). (D) Sucrose gradient fractionation was performed using our engineered HEK293T cell line expressing the 8xHis-Flag-uS7 ribosomal protein. The fractions were used for immunoblotting uS7 with a Flag antibody. An immunoblot of uS3 is used to compare the incorporation of the tagged uS7 protein into ribosomes with that of another unmodified small subunit ribosomal protein. (E) Gene ontology analysis of proteins differentially associated with ribosomes as a function of formaldehyde concentration used for crosslinking. (F) Trajectories of selected protein families in response to formaldehyde doses. When several proteins belong to the same cluster, the mean trajectory (bold line) and the standard deviation (light-colored area) are shown. Otherwise, individual trajectories are shown. (G) Venn diagram showing the total number and overlap of ribosomal core proteins and ribosome-associated proteins identified by our eRAP method and two previous studies^5,19^. (H) Comparison of the top enriched gene ontology categories of our ribosome interactome with those of two previous studies^5,19^. (I) The ribosome interactome comprises various functional protein groups that act at different stages of translation. The number of factors in each group and the identity of its representative factors are indicated.

While optimizing eRAP from uS7-tagged HEK293T cytoplasmic lysates, we noticed that several known RAPs such as initiation factors were not detected, as previously reported in ribosome interactome studies conducted under native conditions^5,6,19^. This finding suggests that weak and/or transient interactions with the ribosome may be lost during the purification process. To overcome this technical issue and also to prevent post-lysis association of spurious proteins, we decided to implement an *in situ* crosslinking step using formaldehyde, a permeable small molecule that efficiently preserves protein complexes by establishing covalent bonds between adjacent lysines and arginines^20^. One risk of using formaldehyde is excessive crosslinking, which can covalently stabilize long-distance and stochastic interactions. Hence, it was critical to identify the optimal formaldehyde concentration to stabilize specific protein interactions while avoiding over-crosslinking. Cells were incubated with different formaldehyde doses (0%, 0.025 %, 0.1 %, 0.37 %, 1 % v/v) and after formaldehyde quenching and cell lysis, samples were incubated with MNase to digest mRNA sequences between multiple translating ribosomes. We selected MNase based on its use in generating rabbit reticulocyte lysates, in which ribosomes remain competent to translate mRNAs. Ribosomes were then tandem-affinity purified using anti-Flag beads coupled with Flag peptide elution, followed by His-Tag magnetic purification coupled with imidazole elution (Figure 1A). Label-free quantitative proteomics was used to identify and quantify RPs and RAPs. In total, 882 different proteins were reliably detected in this dataset (Table S1). Gene ontology (GO) analysis of the detected proteins revealed core ribosomal structural components even in the absence of formaldehyde, while increasing numbers of RAPs were identified as the formaldehyde concentration increased (Figure 1E). Up to 0.37 % formaldehyde, we efficiently recovered proteins involved in translation initiation and elongation, mRNA binding, endoplasmic reticulum, ribosome biogenesis, RNA stability, and RNA processing. Higher formaldehyde doses led to the detection of proteins related to cell cycle and cytoskeleton, indicating the isolation of translation-unrelated proteins, probably through formaldehyde-stabilized long-distance protein-protein bridges.

To gain a more comprehensive understanding of protein recovery as a function of formaldehyde concentrations, we performed a protein clustering analysis, which revealed 16 clusters (Extended Data Figure 1A). Six clusters (1, 3, 4, 6, 14 and 15) exhibited no change with varying formaldehyde doses. In contrast, five clusters (0, 5, 9, 10, and 11) displayed hyperbolic curves. Our interpretation of these results is that long-lived protein-protein interactions have low sensitivity to increasing formaldehyde concentrations, while transitory interactions strongly benefit from increasing formaldehyde concentrations. In agreement with this hypothesis, RPs and late cytoplasmic maturation factors of the 40S subunit (BYSL, LTV1, NOB1, PNO1, RIOK1, TSR1^21^) showed formaldehyde insensitivity (Figure 1F). Recovery of all subunits of eIF3 initiation factor was enhanced by increasing formaldehyde doses, reaching saturation at approximately 0.1 % formaldehyde (Figure 1F). A similar dose-dependent curve was observed for all subunits of the signal recognition particle (SRP) that targets the ribosome-mRNA-nascent polypeptide chain complex to the endoplasmic reticulum (ER) (Figure 1F). Taken together, these data suggest that proteins within a given complex behave similarly towards formaldehyde crosslinking. We then conducted our ribosome interactome experiments using formaldehyde crosslinking, employing 0.1 % formaldehyde to avoid over-crosslinking.

To remove proteins spuriously purified by binding to the beads, we included a negative control corresponding to wild-type cells where uS7 is not tagged (Figure 1A). No proteins were detected co-purifying with untagged uS7 by silver staining (Extended Data Figure 1B). Eluates from wild-type and HF-uS7-tagged cells were compared by label-free quantitative proteomics. This allowed to reliably identify and quantify 884 proteins enriched in eluates from HF-uS7 cells compared to wild-type cells (Table S2). We identified 32 of the 33 small ribosomal subunit proteins and 43 of the 46 large ribosomal subunit proteins. Similar intensities were found for all RPs, with a mean fold enrichment of ∼12 (log_2_ fold change), suggesting that the purified ribosomes are predominantly fully assembled ribosomes. Further supporting this, we detected factors involved in all stages of the translation process, including initiation, elongation, and termination. We also identified ∼800 non-ribosomal proteins, including ∼270 known RAPs and 542 new RAPs. This makes the eRAP strategy the most effective method for identifying proteins specifically associated with ribosomes to date^5,6,16,19,22^. Figure 1G and 1H show a comparative analysis of our interactome with cytoplasmic ribosome interactomes from mouse embryonic stem cells^5^ and neuronal cells^19^. Only 139 proteins are common to all three interactomes, 72 of which are core RPs (Table S3). This limited overlap may be due to differences in cell types and purification methods. GO analysis of the remaining 67 common RAPs revealed that nearly all of them exhibit RNA-binding activity. Our gene clustering analysis showed that 101 of the 139 shared proteins have limited sensitivity to formaldehyde and 29 are highly sensitive. This indicates that most common proteins are involved in long-lived high-affinity protein-protein interactions, while dynamic and labile interactions are underrepresented, confirming the relevance of our formaldehyde crosslinking approach for a more complete picture of the ribosome interactome. We recovered all canonical translation initiation and elongation factors (Tables S2 and S3, Figure 1I), many of which were missing in previous interactomes^5,6,19^. Interestingly, we found EEF1A1, the alpha-1 isoform of the elongation factor 1 complex, but not EEF1A2. However, Tran et al.^19^ found both isoforms. This makes sense when considering the cell types used. EEF1A1 is expressed in the brain and kidney, while EEF1A2 is absent in the kidney, the tissue of origin for HEK293T cells, explaining why it was not detected in our study. GO analysis of the identified 542 novel RAPs revealed enrichment for proteins involved in SRP-dependent protein targeting to ER, translation regulation, proteasome, metabolic and DNA-related processes, among others (Figure 1H). Some of the new RAPs, which comprise distinct functional groups, are indicated in Figure 1I according to the stages of the translation process during which these RAPs are likely to act.

Previous studies^5,6^ showed that nascent peptides do not significantly contribute to MS signals in purified ribosome samples. However, because eRAP is more sensitive than other methods, we investigated nascent peptide incidence in our samples. We compared the read counts of ribosome-protected mRNA fragments (RPF) from HF-uS7-tagged cells with the fold enrichment of proteins identified in the ribosome interactome. This analysis revealed no correlation between RPF counts and protein fold enrichment from eRAP (Extended Data Figure 1C), suggesting that nascent peptides are not a significant source of signal in our samples. Further confirming this result, we also observed no peptide distribution bias toward the N-terminal end in the proteins recovered by eRAP compared to the peptide distribution of the entire cell proteome (Extended Data Figure 1D).

### Differential remodeling of the ribosome interactome by viruses

Viruses rely on host ribosomes to translate their mRNAs and have evolved numerous strategies to hijack cellular translation for their own benefit^10^. Our hypothesis is that virus infection may induce changes in ribosome composition to fine-tune translation of unusual mRNAs such as innate immunity and viral transcripts. A previous study suggested that poliovirus, dengue virus, and Zika virus cause changes in ribosome interactors^16^. However, that study isolated total messenger ribonucleoprotein (mRNP) complexes, rather than purified ribosomes. We believe that eRAP is an ideal method for determining whether viruses can specifically alter the composition of core ribosomes and associated proteins, while excluding other RNA-binding proteins (RBPs) that may co-sediment with polysomes. We chose to analyze two very different viruses: HSV-1 and SINV. HSV-1 replicates in the nucleus and contains a double-stranded DNA genome greater than 150 kilobases that codes for over 80 proteins. It is known to infect humans with a broad cell tropism. To study if the core ribosome and its interactome change during HSV-1 lytic infection, HF-uS7-tagged HEK293T cells were mock-infected or infected with the HSV-1 strain 17 syn^+^ for 8, 12, and 18 hours. To exclude proteins that co-purify non-specifically with ribosomes during the eRAP procedure, we used wild-type HEK293T cells infected with HSV-1 for 12 hours as a control. We identified approximately 650 proteins as RAPs across all time points of infection (Table S4). Some of these proteins were differentially recruited to ribosomes during infection (log□ fold change ≥ 1 or ≤ −1, with a 5% false discovery rate). We detected 30, 68, and 109 viral or cellular proteins that were differentially associated with ribosomes at 8, 12, and 18 hours post-infection (hpi), respectively, when compared to uninfected cells (Figure 2A). HSV-1 infection did not alter the composition of core RPs, and only a few canonical translation factors were more abundantly associated with ribosomes upon infection (colored dark red). HSV-1 also increased the binding to ribosomes of the three components of a complex involved in the UFMylation pathway^23,24^ (colored red). Conversely, HSV-1 infection decreased the binding to ribosomes of several chaperones and heat shock proteins (colored intense blue) and various nuclear proteins (colored turquoise). HERC5, an innate immune protein with broad-spectrum antiviral activity, was also less abundant upon HSV-1 infection (colored green), suggesting a possible virus-induced mechanism to block interferon-mediated antiviral effectors.

**Figure 2.**
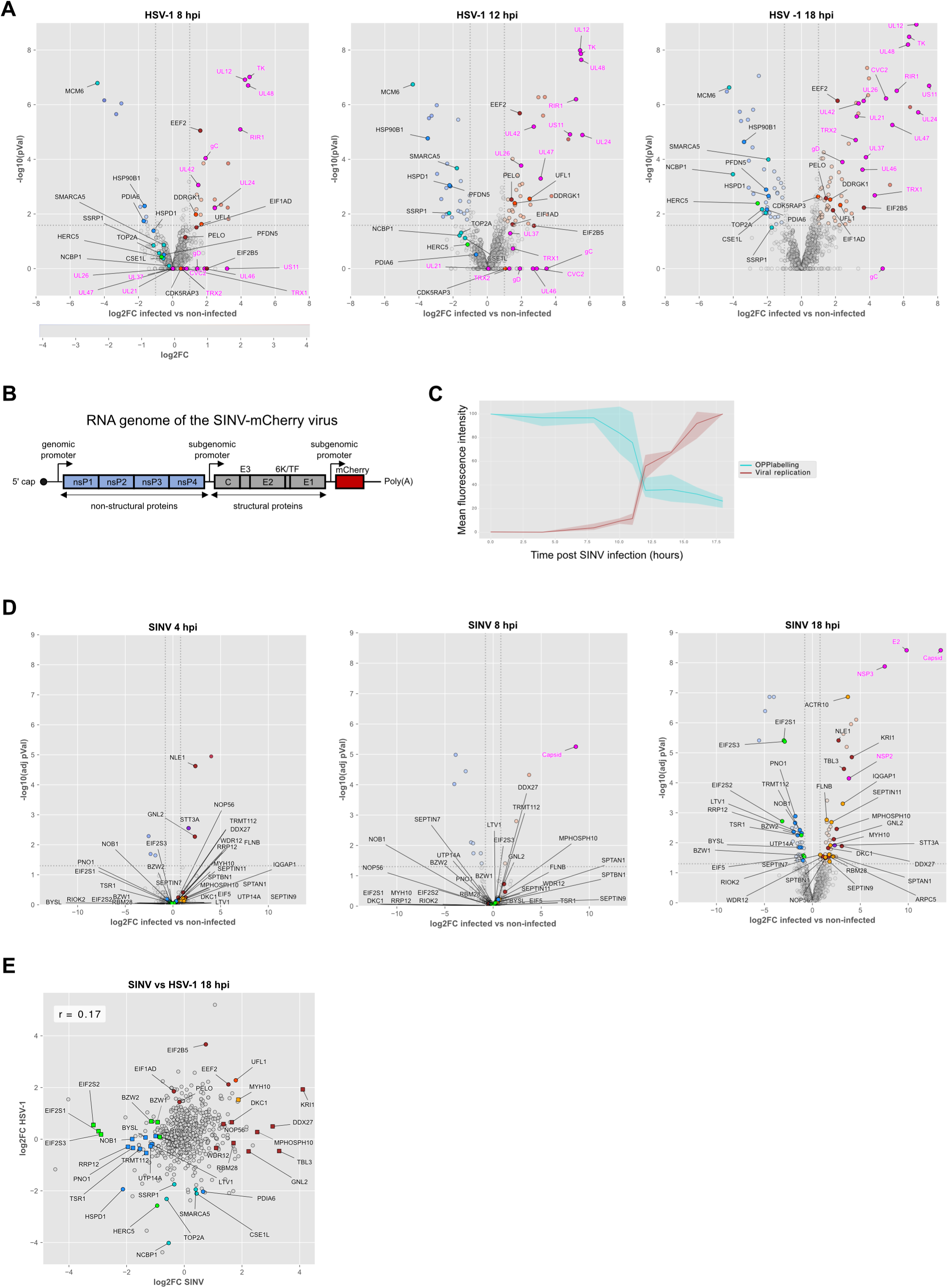
Viruses remodel the ribosome interactome differently. (A) Volcano plots showing changes in the abundance of ribosome-associated proteins at 8, 12, and 18 hours after HSV-1 infection compared to mock-infected cells. Viral proteins are colored fuchsia. Certain ribosome-associated proteins of interest are shown in different colors, as explained in the text. (B) Schematic representation of the genomic structure of the recombinant Sindbis-mCherry virus. (C) Flow cytometry analysis of OPP labeling and mCherry expression in HEK293T cells at various time points following SINV infection. The colored lines represent the means for the two measurements and the colored surfaces represent the 95% confidence interval of the means (n = 3 independent experiments). (D) Volcano plots showing changes in the abundance of ribosome-associated proteins at 4, 8, and 18 hours after SINV infection compared to mock-infected cells. Viral proteins are colored fuchsia. Certain ribosome-associated proteins of interest are shown in different colors, as explained in the text. (E) Scatter plot showing changes in the abundance of ribosome-associated proteins 18 hours after infection with either SINV or HSV-1. The color code is the same as that use in the volcano plots shown in panels A and D. The selected ribosome-associated proteins in cells infected with HSV-1 or SINV are represented by circles and squares, respectively.

The second virus studied was the prototypical alphavirus SINV^25^, an enveloped virus with a positive-sense, single-stranded RNA genome of about 11 kilobases that encodes 10 proteins (Figure 2B). SINV replicates in the cytoplasm of host cells and produces three viral RNAs: 1) a genomic RNA that is translated shortly after viral entry and encodes four non-structural proteins (NSP1-4), which form the replicase complex; 2) a negative-strand RNA that serves as a replication template; and 3) a subgenomic RNA that is transcribed from a duplicated subgenomic promoter and is translated at late times post-infection (starting at 4 hpi) to produce the six structural proteins. To easily track viral spread, we used a previously described chimeric SINV virus expressing the fluorescent mCherry protein from a duplicated subgenomic promoter^26^ (Figure 2B). To determine the optimal infection times for studying the ribosome and its interactome, we first followed viral spread in our HF-uS7-HEK293T cell line. In parallel, we monitored protein synthesis using O-propargyl-puromycin (OPP), a clickable puromycin analog that incorporates into elongating polypeptide chains. Cells were mock-infected or infected with SINV (multiplicity of infection (MOI)=5) for 4 to 18 hours before performing flow cytometry assays (Figure 2C). As early as 12 hpi, SINV infection resulted in a ∼50% decrease in OPP uptake, coinciding with a ∼50% increase in viral spreading. These results are consistent with the expected shutoff of host translation by the virus^26^. Based on the assumption that viral protein synthesis would be low at 4 hpi, dominate at 18 hpi, and be intermediate at 8 hpi, we used eRAP to study the ribosome interactome at these time points (Table S5). Wild-type cells were infected in parallel as controls. Very few viral and cellular proteins were differentially recruited to ribosomes at 4 and 8 hpi (Figure 2D). This number increased to 90 at 18 hpi, 4 of which were viral proteins (colored fuchsia). Several proteins involved in ribosome biogenesis (colored dark red) and several cytoskeletal proteins, including septins (colored orange), exhibited increased binding to ribosomes following SINV infection. Conversely, several factors that act sequentially in 40S subunit maturation^21^ were decreased (colored intense blue). This decrease may reflect SINV-mediated inhibition of 40S biogenesis, but could also arise, at least in part, from the translational shutoff triggered by the virus that restrict synthesis of new RPs. Specific translation initiation factors (colored green) were also decreased. One is eIF2, a heterotrimer that loads the initiator transfer RNA (tRNA) onto the 40S subunit. At 18 hpi, all eIF2 subunits (EIF2S1-3) were strongly depleted. This is likely due to EIF2S1/eIF2α phosphorylation by the dsRNA-activated kinase PKR during alphavirus infection^26^, which blocks eIF2 recycling and subsequent initiation rounds. A second factor is eIF5, which interacts with EIF2S2 and promotes hydrolysis of eIF2-bound GTP upon start codon recognition^27^. Additionally, eIF5 favors initiation in weak contexts^28^. This activity is counteracted by BZW1 and BZW2, thereby increasing start codon stringency^29^. Interestingly, BZW1 and BZW2 exhibited decreased binding to ribosomes at 18 hpi. These findings suggest that SINV infection disrupts the availability of the ternary complex as well as the regulatory balance governing start codon selection. Finally, the cap-binding initiation factors eIF4E, eIF4G, and the helicase eIF4A were slightly reduced, though not statistically significant. Overall, our results imply that SINV infection alters translation initiation by decreasing the association of several initiation factors and regulators of start codon selection with the 40S subunit.

Next, we compared the changes in the ribosome interactomes of cells infected with SINV and HSV-1. This analysis revealed no global correlation between the two conditions (Figure 2E). The core ribosomal proteins remained largely unchanged in both infections, whereas most RAP alterations were virus-specific. This suggests that, despite relying on the same host ribosomes, these two unrelated viruses have evolved distinct strategies for utilizing the host cell translation machinery to optimize viral protein synthesis and disable host antiviral responses. These findings also suggest that the observed RAP alterations do not result from a generic host cell stress response to viral infection.

### Remodeling of the polysome interactome by SINV

Because SINV shuts off host mRNA translation, likely resulting in the accumulation of non-translating ribosomes, we set out to determine the composition and interactome of polysomes in infected cells, which should primarily correspond to ribosomes engaged in translating viral mRNAs. For this, wild-type and HF-uS7-tagged HEK293T cells were mock-infected or infected with SINV for 12 hours. We chose a 12-hour infection time in order to preserve an adequate quantity of polysomes for eRAP (Figure 2C). After formaldehyde crosslinking, cell lysates were layered on top of sucrose gradients and ultracentrifuged to separate ribosomal subunits, 80S ribosomes, and polysomes. The polysomal fractions were pooled and treated with MNase before tandem affinity purification and quantitative proteomic analysis (Figure 3A). Following viral infection, we identified 67 host proteins that were differentially associated with polysomes (Table S6). The association decreased for 13 of these proteins upon infection, including several immune response proteins such as the kinase PKR, also known as EIF2AK2 (Figure 3B). Among the 54 factors enriched in the polysomes of infected cells compared to non-infected ones, several are involved in translation and translation quality control (Figure 3B, colored turquoise). DENR and MCTS1 form a heterodimer that acts as a non-canonical translation initiation factor^30^.

**Figure 3.**
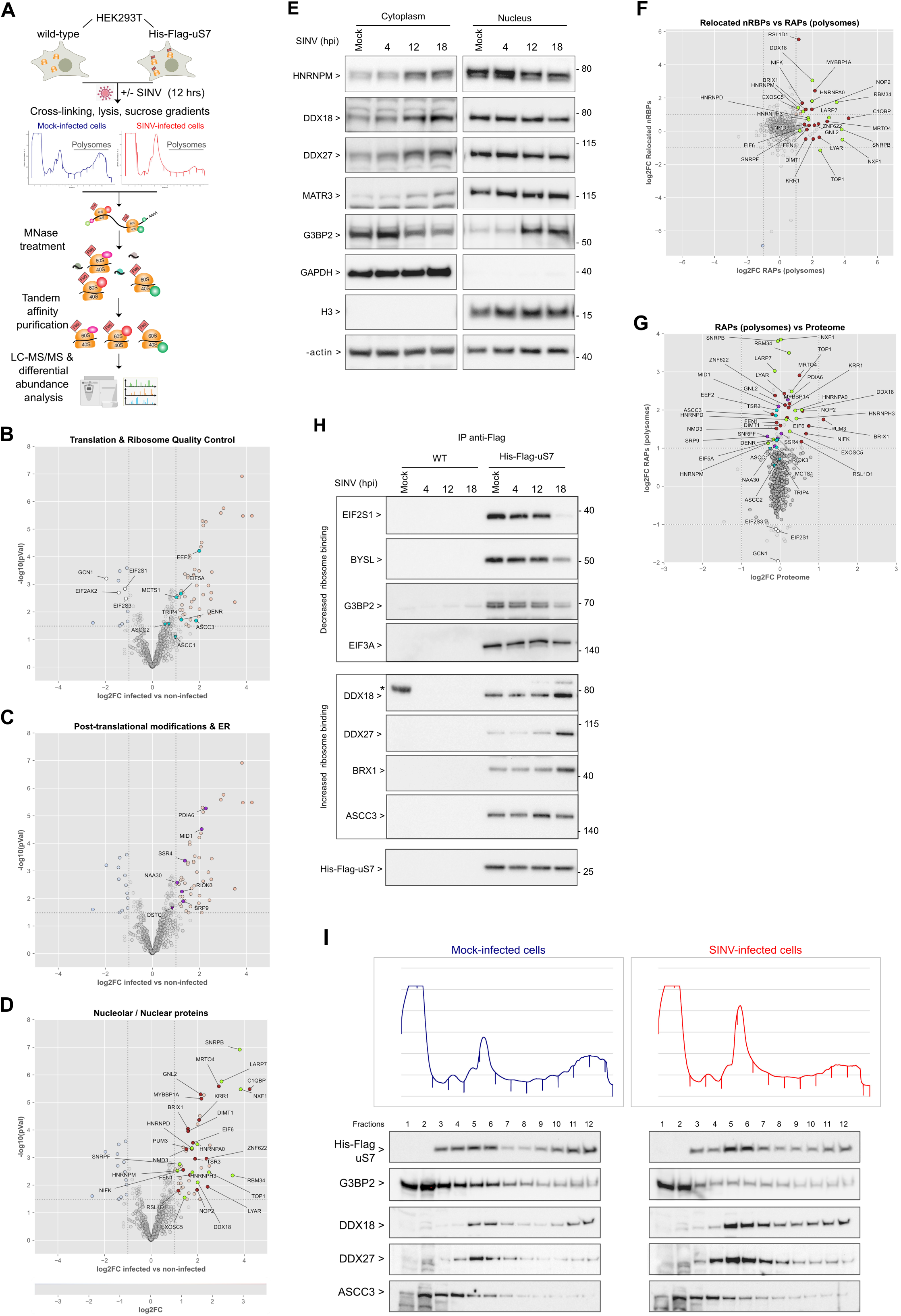
The polysome interactome is remodeled by SINV. (A) Schematic of the eRAP method implemented to determine the polysome interactome. After crosslinking and lysis of the cells, the polysomes were separated using sucrose density gradient centrifugation. The pooled polysomal fractions were then subjected to eRAP. (B) Volcano plot showing changes in the abundance of proteins associated with polysomes 12 hours after SINV infection compared to mock-infected cells. Proteins involved in translation and ribosome quality control that have higher or lower abundance are highlighted in turquoise and white, respectively. Viral proteins are not shown on the volcano plot. (C) The same volcano plot as in (B). Proteins involved in co/post-translational modifications and protein folding are highlighted in purple. (D) The same volcano plot as in (B). Various proteins that are primarily localized in the nucleolus or nucleus are highlighted in different colors, as explained in the text. (E) The cytoplasm and nucleus of mock-infected cells and cells infected with SINV for 4, 12, or 18 hours were separated and used for different immunoblots. Immunoblots for GAPDH and histone H3 are used as quality controls for the cytoplasmic and nuclear fractions, respectively. The immunoblot for β-actin serves as a loading control. (F) Scatterplot showing the relationship between changes in the abundance of polysome-associated proteins and the nucleo-cytoplasmic redistribution of nuclear RBPs in response to SINV infection. Kamel et al.^34^ calculated the protein redistribution index as the (log_2_) ratio of protein intensity in the cytoplasmic fraction to the nuclear fraction. The highlighted proteins are color-coded as in Figure 3D. (G) Scatterplot showing the relationship between changes in total protein abundance and changes in polysome association following SINV infection. The highlighted proteins are color-coded as in Figures 3B-D. (H) Flag immunoprecipitations from parental cells and cells expressing His-Flag-uS7 that were either left uninfected or infected with SINV for 4, 12, or 18 hours. Different immunoblots of the eluates are shown. The His-Flag-uS7 immunoblot is used as a control for equal pull-down efficiency. Of note, the asterisk in the first lane of the DDX18 immunoblot indicates a nonspecific signal caused by the molecular weight marker. (I) HEK293T cells expressing His-Flag-uS7 were either mock-infected or infected with SINV for 12 hours. Then, the different ribosomal populations were separated by sucrose density gradient centrifugation (upper panels). The resulting fractions were used for different immunoblots, as indicated.

eIF5A was initially identified as an initiation factor; however, subsequent studies revealed its role in promoting translation elongation and termination, especially when ribosomes stall at specific amino acid sequences^31^. ASCC3 is a helicase in a complex that rescues collided ribosomes^32,33^. The other three components of this complex (ASCC1, ASCC2, and TRIP4) were also enriched, although they did not pass our stringent cutoff.

Another group of enriched proteins is involved in co- and post-translational modifications, as well as protein folding (Figure 3C, colored purple). RIOK3 and MID1 have kinase and E3 ubiquitin ligase activities, respectively. NAA30 catalyzes N-terminal acetylation. Other ER-connected proteins were also identified: SRP9, a subunit of the SRP complex that targets secretory and membrane proteins to the ER; SSR4, a component of the translocon associated protein (TRAP) complex that stimulates the cotranslational translocation of nascent ER proteins; PDIA6, which exhibits isomerase and chaperone activities in the ER. Additionally, OSTC, a subunit of the STT3A-containing oligosaccharyltransferase (OST-A) complex that catalyzes cotranslational N-linked glycosylation in the ER lumen, was enriched, though not statistically significant. The catalytic subunit STT3A, however, was significantly enriched in the total ribosome interactome (Figure 2D, colored purple).

The last group of factors enriched in polysomes during SINV infection includes proteins involved in ribosome biogenesis (Figure 3D, colored dark red). Others are RBPs, including heterogeneous nuclear ribonucleoproteins (hnRNPs), as well as proteins involved in mRNA processing, splicing, and export (Figure 3D, colored green). Many of these proteins, which have also been identified in the total ribosome interactome (Table S5), are primarily localized in the nucleolus or nucleus. This finding was intriguing for polysome-associated proteins. To further validate this, we performed nuclear/cytoplasmic fractionation on mock- and SINV-infected cells. We then detected some of the nucleolar/nuclear RAPs identified in either of our two interactomes by immunoblotting. During SINV infection, we observed partial relocation of HNRNPM, DDX18, DDX27, and MATR3 to the cytoplasm (Figure 3E). Conversely, G3BP2 redistributed to the nucleus during infection. These bidirectional changes in RBP subcellular localization, coupled with the absence of changes in the control proteins GAPDH, histone H3, and β-actin, suggest that the nuclear envelope remains intact during infection. Consistent with our findings, it has been recently reported that SINV induces the redistribution of numerous nuclear RBPs into the cytoplasm^34^. Notably, we identified many of these RBPs in our polysome interactome (Figure 3F).

To exclude the possibility that the aforementioned proteins interact with polysomes simply due to higher levels in infected cells, we analyzed the proteome of mock- and SINV-infected cells using quantitative proteomics. Then, we compared changes in theproteome and polysome interactome after SINV infection. Our results showed that infection does not significantly alter the total levels of these polysome-associated proteins (Figure 3G, Table S7), indicating that the observed changes in polysome-association are actively regulated and not passive events.

To validate our eRAP results, we performed Flag-tagged ribosome immunopurification and immunoblotting of specific RAPs identified in the polysome or total ribosome interactomes (Figure 3H). To align with the time points used for the interactomes, we infected or mock-infected wild-type and HF-uS7-tagged HEK293T cells with SINV for 4, 12, or 18 hours. All of the selected RAPs exhibited behaviors consistent with the proteomics data. Notably, ASCC3 was again more abundantly immunoprecipitated at 12 hpi than at 18 hpi. We also performed sucrose gradient fractionation coupled with immunoblotting on both mock-infected and 12-hour-infected cells. A relative comparison of the co-sedimentation profiles revealed that DDX18 and DDX27 are more abundant and G3BP2 is less abundant in all ribosomal fractions of infected cells (Figure 3I). ASCC3 was only more abundant in polysomes. These results corroborate our proteomics data and strongly suggest that the identified factors are bona fide RAPs that alter their association with ribosomes in response to SINV infection.

### SINV infection triggers distinct regulatory layers of RNA-binding proteins on mRNAs and ribosomes

Since many RAPs from our datasets are also known RBPs, we investigated whether viral infection affects the ribosome interactome and RBPome similarly. First, we analyzed published RBPome datasets obtained by RNA interactome capture (RIC) of poly(A) mRNAs from cells infected with SINV^26^, which allow to measure changes in RNA-binding activity of RBPs upon infection. We compared changes in RBP-binding activity at 18 hpi with our global ribosome interactome dataset obtained at the same timepoint. This comparison revealed a lack of correlation between the two datasets, with only a few RBPs displaying concordant changes (Figure 4A). Several ribosome-processing factors, including BYSL, NCL, RRP12, UTP14A, and XRCC6, were significantly down-regulated, while EEF2 and LARP7 were significantly up-regulated in both the ribosome interactome and RBPome (colored yellow). In contrast, many RBPs showed opposite trends in the two datasets (colored turquoise), while most remained unchanged or underwent selective regulation in the RBPome (colored green) or ribosome interactome (colored orange). This lack of global correlation could be explained by the translational shutoff induced by infection, which restricts translation of most cellular mRNAs in favor of viral RNA translation (Figure 2C). To further complement this analysis, we integrated published viral RNA interactome capture (vRIC) datasets that identify cellular and viral RBPs directly bound to all SINV RNAs^34^, independently of whether these RNAs are engaged in translation. To pinpoint candidate cellular RBPs with a potential role in viral RNA translation, we examined the distribution of RBPs identified by vRIC across changes in polysome association at 12 hpi, a condition in which polysomal ribosomes predominantly translate viral RNAs. The RBPs identified in the vRIC dataset were distributed evenly and broadly among the proteins exhibiting increased or decreased association with polysomes upon infection (Figure 4B, burgundy). This suggests that polysome enrichment cannot be predicted solely by viral RNA binding. Notably, members of the DEAD-box and DExH-box RNA helicase families were preferentially skewed toward increased polysome association during infection (Figure 4B, green), consistent with a potential role in controlling viral RNA translation (although most did not meet the predefined cutoffs for fold-change and statistical significance, with the exception of DDX18). Additionally, ASCC2 and ASCC3 (colored turquoise) bind viral RNAs and display increased association with polysomes during infection, thus emerging as additional candidate factors linking viral RNA binding to translational engagement.

**Figure 4.**
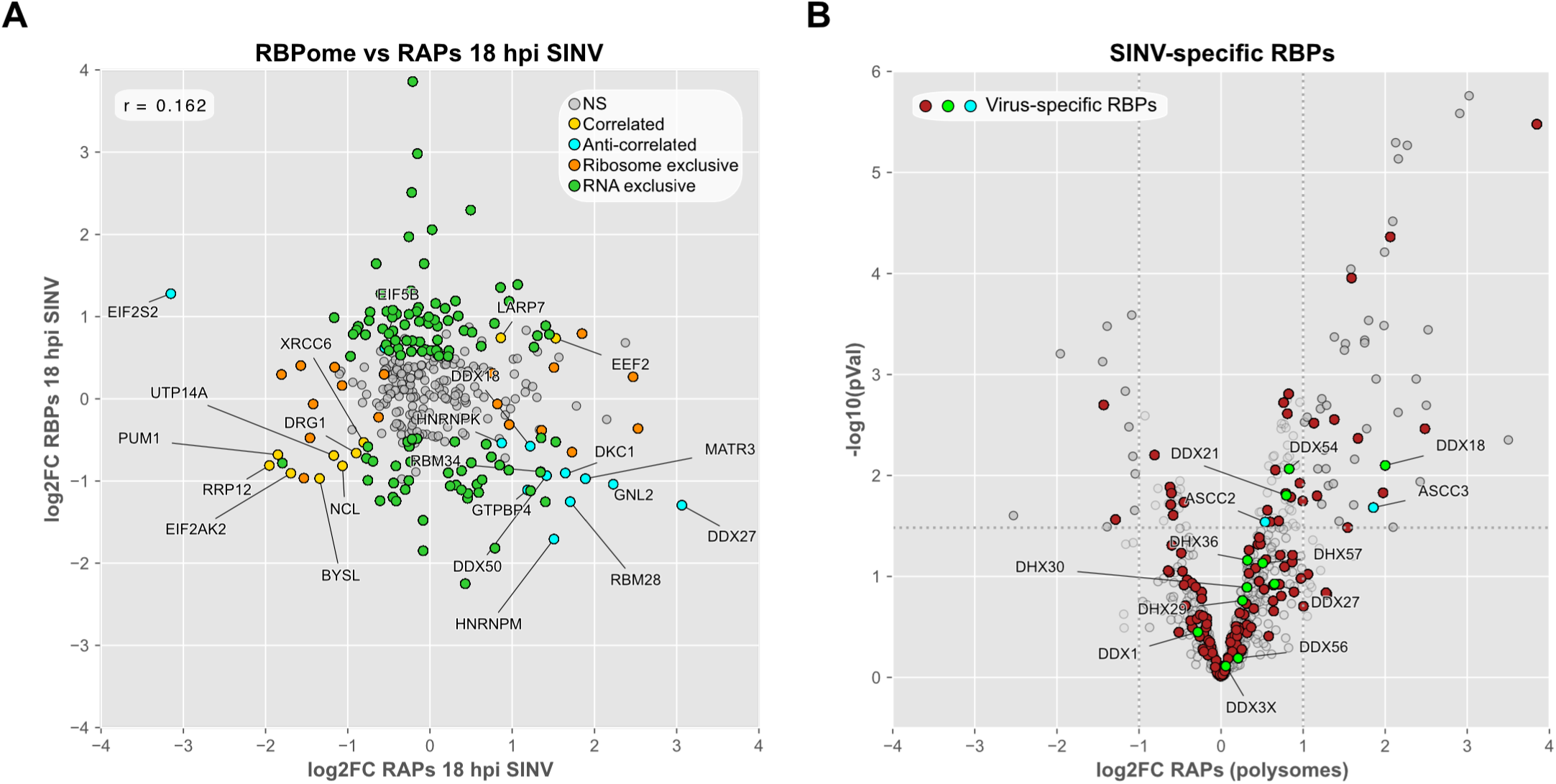
(A) Scatter plot showing the relationship between changes in RNA-binding protein (RBP) recruitment to poly(A)-tailed mRNAs at 18 hpi with SINV (y-axis) and changes in their recruitment to ribosomes (x-axis). (B) Volcano plot showing changes in polysome-associated proteins at 12 hpi with SINV. RBPs previously identified as specifically binding to viral RNAs are colored burgundy. DEAD-box and DExH-box helicases, as well as ASCC2 and ASCC3, which also bind viral RNAs, are highlighted in green and turquoise, respectively. Viral proteins are not shown on the volcano plot.

Taken together, these results imply that changes in RNA-binding activity and ribosome association are largely uncoupled during SINV infection. However, combining polysome eRAP profiling with viral RNA interactome data reveals candidate RBPs associated with translating viral RNAs.

### SINV remodeling of the ribosome interactome is functionally important

To determine if proteins enriched in the ribosome interactome are necessary for SINV infection, we examined the effects of suppressing some of them using CRISPR/Cas9 gene editing. Gene disruption was performed in bulk in HEK293T cells, which were subsequently infected with SINV-mCherry at a low MOI for 24 hours. We then measured the number of mCherry-positive cells by flow cytometry analysis (Figure 5A). The first group we tested comprises RAPs that localize to the nucleolus or nucleus in uninfected cells but relocate to the cytoplasm following SINV infection (Figure 3E). We found that deleting the DEAD-box RNA helicases DDX18 and DDX27 reduced SINV infection (Figure 5B). However, knocking out the ribonucleoproteins HNRNPM and MATR3 had little impact on the number of mCherry-positive cells (Figures 5B). Deleting the endonuclease FEN1 did not affect SINV propagation either (Figure 5B). Although these RAPs have been reported to promote or inhibit viral replication via various mechanisms^35–40^, our results show that only two of them are critical host factors for SINV.

**Figure 5.**
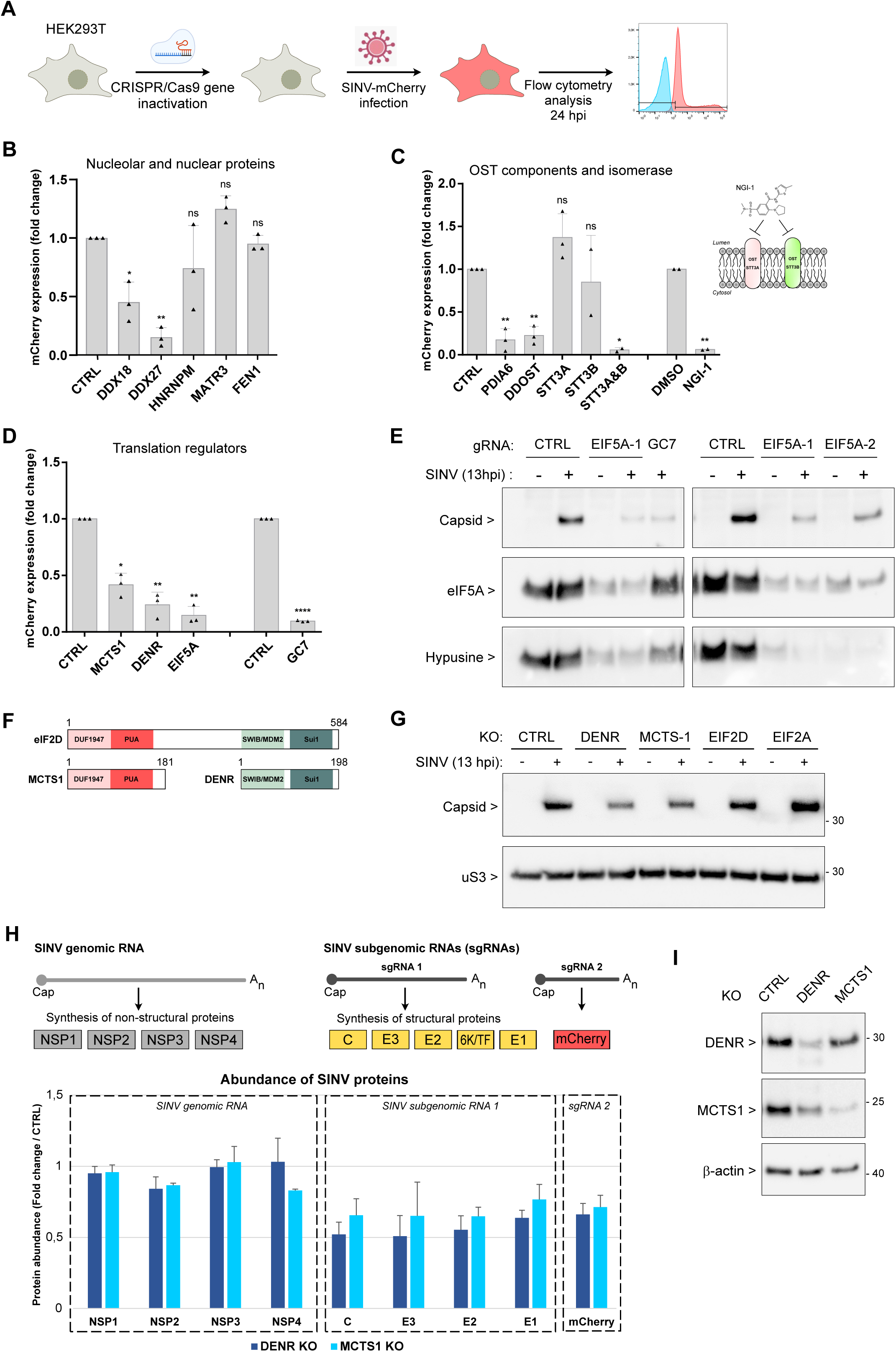
(A) Experimental strategy for evaluating the importance of ribosome-associated proteins in SINV replication. (B) mCherry fluorescence is used to quantify SINV replication in cells invalidated for genes encoding various ribosome-associated proteins primarily localized in the nucleolus or nucleus of uninfected cells. The bar plot shows the mean of three biological replicates, with each replicate consisting of three to four technical replicates. The error bars represent the standard error of the mean. Two-tailed paired t-tests were performed to compare the mCherry signal in individual knockout cells to the signal in control knockout cells. *p < 0.05, **p < 0.01, ***p < 0.001, and ****p < 0.0001. “ns” indicates not significant. (C) Bar plot showing the quantification of mCherry fluorescence in cells transfected with the indicated sgRNAs to edit genes encoding some OST components and PDIA6. The bar plot also shows results of cells infected with SINV and treated simultaneously with 5 µM of NGI-1 or DMSO for 24 hours. The bar plot represents the mean of two or three biological replicates, with each replicate consisting of three to four technical replicates. The error bars and statistical tests are as described in Panel B. The mCherry signal in NGI-1-treated cells was compared to the signal in vehicle-treated cells. (D) Bar plot showing the quantification of mCherry fluorescence in cells transfected with the indicated sgRNAs to edit genes that encode specific translation factors. In addition, results of cells infected with SINV and treated simultaneously with 25 µM of GC7 or vehicle for 24 hours are shown. The bar plot represents the mean of three biological replicates, with each replicate consisting of three to four technical replicates. The error bars and statistical tests are as described in Panel B. The mCherry signal in GC7-treated cells was compared to the signal in untreated cells. (E) Immunoblot analysis of SINV capsid protein levels in EIF5A knockout cells and GC7-treated cells. The cells were infected at a high MOI for 13 hours and, where indicated, treated simultaneously with 25 µM of GC7. Immunoblots of total eIF5A and hypusinated eIF5A are also shown. (F) Schematic representation of the human eIF2D, MCTS1, and DENR proteins, showing their homologous domains. (G) Immunoblot analysis of SINV capsid protein levels in cells lacking DENR, MCTS1, eIF2D, or eIF2A. The cells were infected at a high MOI for 13 hours. An immunoblot of uS3 serves as a loading control. (H) Quantification by mass spectrometry of all SINV proteins synthesized from genomic and subgenomic RNAs in cells lacking DENR or MCTS1, compared to control cells. The cells were infected at a high MOI for 13 hours. (I) Immunoblots showing the efficiency of knocking out DENR and MCTS1 in the cell lysates used for the mass spectrometry analysis shown in Panel H. The β-actin immunoblot serves as a loading control.

Next, we sought to determine the importance of the OST complex for SINV. Previous studies have shown that it is essential for flaviviruses^41,42^. Mammalian cells have two OST complex isoforms. One contains the STT3A catalytic subunit, and the other contains STT3B. These two OST isoforms possess common and distinct accessory subunits. Complete cotranslational N-linked glycosylation of polypeptides requires the cooperation of both isoforms^43,44^. Only STT3A was recovered in our interactomes. We deleted either STT3A or STT3B, or both, to completely prevent N-glycosylation since STT3B can partially rescue STT3A^43^. We also deleted DDOST, a subunit shared by both OST isoforms. We found that suppressing just one catalytic subunit had little to no impact on the number of mCherry-positive cells (Figure 5C). However, co-suppressing STT3A and STT3B almost completely prevented infection. Deleting DDOST also had a significant inhibitory effect. Similar results were obtained using a second set of single-guide RNAs (sgRNAs) targeting these OST genes (Extended Data Figure 5A). Furthermore, deleting these genes decreased the production and infectivity of progeny virions (Extended Data Figures 5B-C). To further study how SINV relies on the OST complex, we tested the effect of NGI-1 treatment on viral replication. NGI-1 inhibits both OST isoforms and has been shown to block the replication of several flaviviruses, but not the alphaviruses Chikungunya (CHIKV) and Venezuelan equine encephalitis (VEEV)^45^. We infected and treated HEK293T cells simultaneously with NGI-1 (5 µM) or the vehicle for 24 hours. We found that NGI-1 treatment strongly inhibited SINV infection (Figure 5C). This suggests that SINV and the two other alphaviruses, CHIKV and VEEV, may differ in their dependency on the OST complex for cotranslational glycosylation of viral proteins. We also investigated whether SINV requires the ER isomerase PDIA6 and found that suppressing it significantly reduced the number of mCherry-positive cells (Figure 5C). PDIA6 is known to restrict the activation of the unfolded protein response (UPR) in case of ER stress, by binding to the UPR effector IRE1α in the ER lumen^46^. We therefore examined whether SINV induces ER stress using IRE1α-mediated expression of spliced XBP1 as a readout. We detected no spliced XBP1 in PDIA6-deficient cells infected with SINV (Extended Data Figure 5D), which indicates an absence of major ER stress. However, PDIA6 may be necessary for the proper folding of SINV glycoproteins, as was previously demonstrated for the E1 glycoprotein of VEEV^47^.

Next, we investigated whether SINV required specific translation factors enriched in the polysome interactome, including eIF5A and the heterodimer DENR-MCTS1 (Figure 3B). We found that eIF5A, DENR, and MCTS1 are important for SINV propagation (Figure 5D, Extended Data Figure 5E). To date, eIF5A is the only protein identified as undergoing hypusination. This post-translational modification converts a lysine residue into hypusine, a modified lysine with a 4-aminobutyl group from the polyamine spermidine^48^. Hypusination of eIF5A is essential to its activity^49^ and can be inhibited by N^1^-guanyl-1,7-diaminoheptane (GC7)^50^. We tested the importance of hypusination for SINV by infecting HEK293T cells at a low MOI and simultaneously treating them with GC7 or the vehicle for 24 hours. GC7 treatment strongly inhibited SINV infection (Figure 5D). We also performed immunoblots that showed that suppressing eIF5A or treating with GC7 decreased the amounts of the viral capsid protein (Figure 5E). Overall, our findings indicate that hypusine-modified eIF5A is necessary for SINV. One might expect that the enrichment of eIF5A in the polysome interactome would facilitate viral RNA translation elongation, particularly when ribosomes encounter difficult-to-translate sequences. Interestingly, opposing trajectories were observed between DENR-MCTS1 and eIF2 with respect to ribosome association during infection. DENR-MCTS1 levels increased in the polysome interactome, while eIF2 levels decreased in the total ribosome interactome (Figure 2D, 18 hpi time point). eIF2 is the primary factor that delivers the initiator tRNA in the canonical translation initiation pathway. In contrast, eIF2A, eIF2D, and DENR-MCTS1 act as alternative factors that deliver the initiator tRNA in a condition-or transcript-selective manner^30,51,52^. MCTS1 and DENR are homologous to the N-and C-terminal regions of eIF2D, respectively (Figure 5F). A previous work^53^ suggested that SINV employs an alternative pathway to initiate viral translation, thereby overcoming the inhibition of translation initiation caused by eIF2 phosphorylation. We then examined whether SINV requires the aforementioned alternative initiation factors. We infected different knockout cells with SINV and assessed capsid protein synthesis by immunoblotting. No changes in capsid amounts were observed in eIF2D- or eIF2A-deleted cells (Figure 5G). However, suppressing DENR or MCTS1 decreased capsid levels (Figure 5G), consistent with our flow cytometry results (Figure 5D). These findings imply that the DENR-MCTS1 dimer is specifically required for SINV and cannot be substituted by eIF2D or eIF2A. Consistent with our results, one study demonstrated that neither eIF2D nor eIF2A is necessary for SINV subgenomic mRNA translation when eIF2 is inactive^54^. However, that study did not examine the impact of DENR-MCTS1. Our findings suggest that DENR-MCTS1 may be the missing link. Next, we examined the effects of DENR-MCTS1 loss on translation of SINV genomic and subgenomic RNAs. We performed quantitative proteomics on infected knockout cells. Suppressing DENR or MCTS1 did not significantly alter the production of non-structural proteins from genomic RNA (Figure 5H, Table S8). However, loss of DENR or MCTS1 decreased the production of structural proteins from subgenomic RNA and of the mCherry protein from a second subgenomic RNA that shares the same 5’UTR and 3’UTR (Figure 5H, Table S8). Gene knockout efficiency was controlled by immunoblotting, which showed that suppressing one dimer component compromises the stability of the other (Figure 5I). This supports the idea that DENR and MCTS1 exist as a complex. Overall, our results suggest a novel role for DENR-MCTS1 in SINV subgenomic RNA translation. This finding further confirms the results of previous *in vitro* experiments showing that DENR-MCTS1 can load an initiator tRNA onto a reporter mRNA bearing the 5’UTR of the SINV subgenomic RNA ^55^.

### A novel non-canonical role for ASCC3 in SINV-infected cells

Among the changes in polysome composition upon infection with SINV, we found that ASCC3 was enriched in polysomes of infected cells (Figure 3B). Furthermore, vRIC experiments revealed that ASCC3 binds directly to viral RNAs (Figure 4B). The other components of the ASCC3 complex, including ASCC1, ASCC2, and TRIP4, were also enriched in polysomes, though not at a statistically significant level. These proteins are part of a complex known as the activating signal cointegrator 1 complex (ASCC) or the ribosome-associated quality control-trigger (RQT) complex. The RQT complex was so named because of its role in resolving ribosome collisions ^32,33^. To determine whether SINV co-opts the RQT complex, we tested the effect of suppressing each of its components using our flow cytometry assay. We found that loss of ASCC3 reduced the number of mCherry-positive cells (Figure 6A). Deletion of ASCC2 and ASCC1 had a moderate inhibitory effect, while loss of TRIP4 had no effect (Figure 6A). Immunoblots performed to verify knockout efficiency revealed that ASCC3 deletion led to decreased levels of the other components, supporting their interaction (Figure 6B). Loss of ASCC2 slightly destabilized ASCC3 and TRIP4, while loss of ASCC1 and TRIP4 had no effect on the other components. This likely explain the results of the SINV spreading assay: a strong phenotype when ASCC3 is suppressed, and little to no phenotype when the other subunits are suppressed. Additionally, deleting ASCC3 decreased the production of progeny virions (Extended Data Figures 6A). Ribosomal stalling and collisions are known to induce ribosome-associated quality control (RQC) in order to degrade truncated nascent proteins^56,57^. Structural studies have shown that collided di-ribosomes adopt a specific conformation, with the 40S-40S interface serving as a ubiquitin signaling platform^58,59^. ZNF598 catalyzes the poly- and mono-ubiquitination of the 40S RPs uS10 and eS10, respectively. The uS10 poly-ubiquitination is essential for the subsequent recruitment of the RQT complex^59^. ASCC2 recognizes the poly-ubiquitin chains on uS10, and the helicase ASCC3 dissociates the leading stalled ribosome, enabling the lagging collided ribosome to resume translation^32,33^. In addition to ZNF598-catalyzed marks on collided ribosomes, RNF10 mono-ubiquitinates uS3 and uS5 on collided and arrested ribosomes during translation initiation or elongation^60,61^. To evaluate whether SINV requires the RQT complex to resolve ribosome collision issues, we analyzed ubiquitin marks by immunoblotting. As a positive control, we induced ribosome collisions by treating mock- and SINV-infected cells with a moderate concentration of anisomycin. As expected, levels of ubiquitinated eS10 and uS10 increased in lysates from anisomycin-treated mock-infected cells compared to untreated cells (Figure 6C). These marks were not detected in lysates from SINV-infected cells, regardless of anisomycin treatment. This indicates that ribosome collisions do not occur at detectable levels following infection, even under conditions that facilitate their formation. Conversely, levels of ubiquitinated uS3 and uS5 increased in lysates from SINV-infected cells compared to mock-infected cells indicating significant levels of stalled ribosomes induced by infection (Figure 6C). Additionally, we investigated the impact of suppressing PKR on ribosome collisions following viral infection. Since depleting PKR decreases EIF2S1 phosphorylation in infected cells (Extended Data Figure 6B), it should permit host mRNA translation to some extent during viral infection. We observed slightly increased levels of ubiquitinated eS10 in PKR-deficient cells treated with anisomycin compared to control cells treated the same way (Figure 6D), suggesting that more collisions occur. However, infecting PKR-deficient cells with SINV did not increase ubiquitinated eS10 levels (Figure 6D). Overall, our results indicated that SINV infection leads to increased levels of RNF10-dependent stalling marks, but not ZNF598-dependent collision marks.

**Figure 6.**
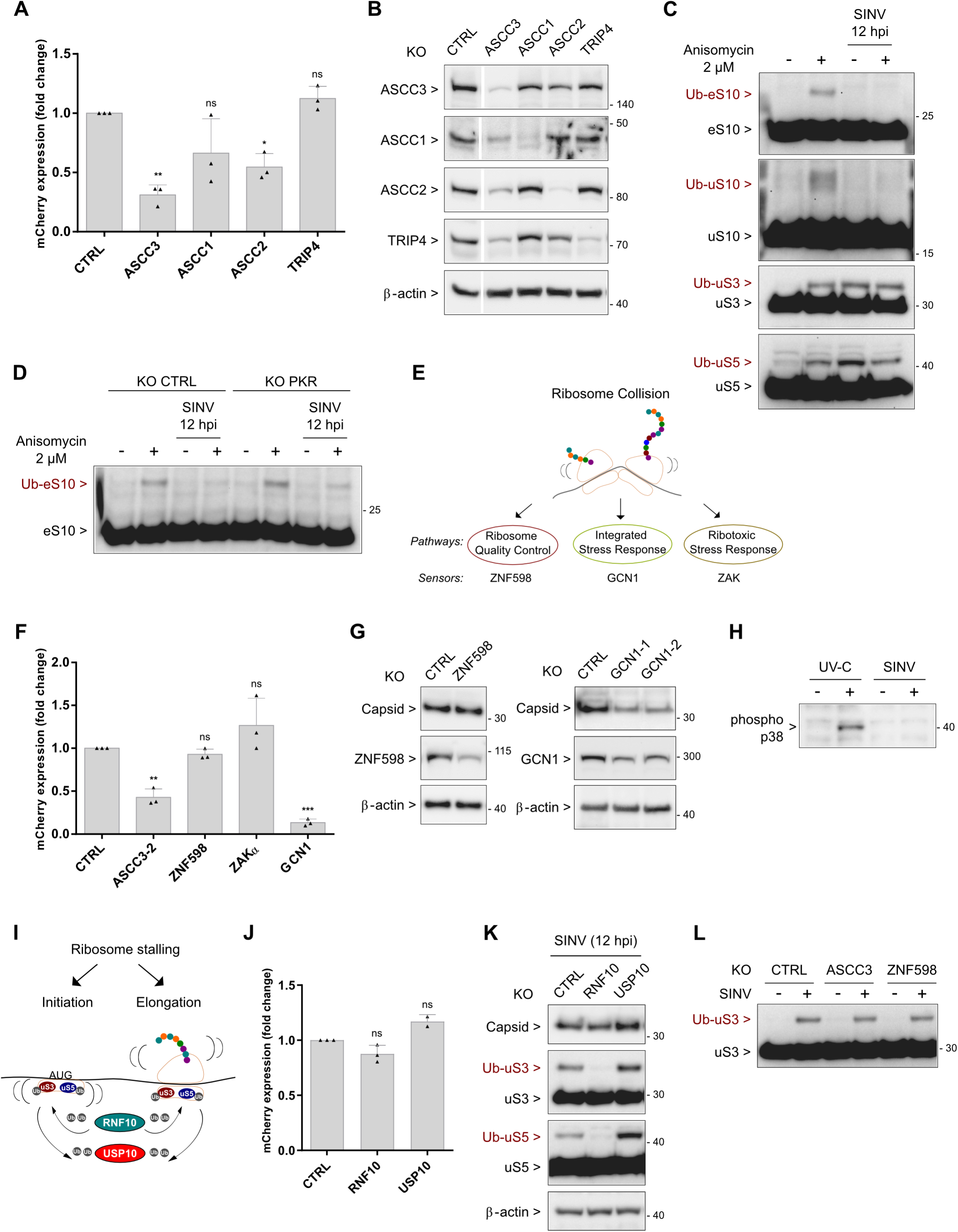
(A) Measurement of mCherry fluorescence serves as an indicator of SINV replication in cells that were transfected with the indicated sgRNA to edit genes encoding the different components of the ribosome-associated quality control-trigger (RQT) complex. The cells were infected at a low MOI for 24 hours. The bar plot shows the mean of three biological replicates, with each replicate consisting of three to four technical replicates. The error bars represent the standard error of the mean. Two-tailed paired t-tests were performed to compare the mCherry signal in individual knockout cells to the signal in control knockout cells. *p < 0.05, **p < 0.01, ***p < 0.001, and ****p < 0.0001. “ns” indicates not significant. (B) Immunoblots showing the efficiency with which each subunit of the RQT complex was knocked out. The β-actin immunoblot serves as a loading control. (C) Cells were either uninfected or infected with SINV at a high MOI for 12 hours. Before collection, the cells were left untreated or treated with 2 µM of anisomycin for 15 minutes, and lysed in RIPA buffer supplemented with 10 mM N-Ethylmaleimide. Cell lysates were used for immunoblotting with antibodies against the ribosomal proteins eS10, uS10, uS3, and uS5. The ubiquitin-modified ribosomal proteins are indicated on the blots as brown colors. (D) Cells were transfected with either a control sgRNA or one targeting PKR. Then, the cells were infected, treated with anisomycin, and lysed as described in panel C. An immunoblot against eS10 is shown, with the ubiquitinated eS10 indicated. (E) Schematic representation of the signaling pathways potentially activated in response to ribosome collisions, along with their corresponding collided ribosome sensors. (F) A bar plot showing the quantification of mCherry fluorescence in cells transfected with the indicated sgRNAs for gene editing and then infected with SINV at a low MOI for 24 hours. The bar plot represents the mean of three biological replicates, each consisting of three to four technical replicates. Error bars and statistical tests are as described in Panel A. (G) Immunoblot analysis of SINV capsid protein levels in cells transfected with *ZNF598*- or *GCN1*-targeting sgRNAs. Two different sgRNAs were tested for *GCN1* gene editing. The cells were infected at a high MOI for 13 hours. The immunoblots below show the efficiency of *ZNF598* or *GCN1* gene knockout, with the β-actin immunoblots serving as loading controls. (H) Immunoblot analysis of ZAKα-mediated p38 phosphorylation in cells infected with SINV at a high MOI for 13 hours. UV-C treatment was used as a positive control for phosphorylated p38. (I) Schematic representation of the ubiquitination and de-ubiquitination of uS3 and uS5 by RNF10 and USP10, respectively, in response to ribosome stalling during translation initiation and elongation. (J) Bar plot showing mCherry fluorescence quantification in cells transfected with the indicated sgRNAs and subsequently infected with SINV at a low MOI for 24 hours. This bar plot represents the mean of two or three biological replicates, with each replicate consisting of three to four technical replicates. The error bars and statistical tests are as described in Panel A. (K) Immunoblot analysis of SINV capsid protein levels in cells transfected with either *RNF10*- or *USP10*-targeting sgRNAs. The cells were infected at a high MOI for 12 hours. Immunoblots against uS3 and uS5 validate the effectiveness of the RNF10 and USP10 knockouts by revealing decreased and increased levels of the uS3/uS5 ubiquitinated forms, respectively. The β-actin immunoblot serves as a loading control. (L) Immunoblot analysis of uS3 ubiquitination in cells transfected with sgRNAs targeting either *ASCC3* or *ZNF598*. The cells were infected at a high MOI for 12 hours.

RQC with ZNF598 as the sensor is not the only surveillance mechanism triggered by ribosome collisions. Under stress conditions that cause extensive ribosome collisions, cells activate either the integrated stress response (ISR) or the ribotoxic stress response (RSR)^62,63^ (Figure 6E). In the ISR pathway, colliding ribosomes are sensed by GCN1, which interacts with the leading stalled ribosome and the lagging collided ribosome^64^. GCN1 then recruits GCN2, another kinase that phosphorylates EIF2S1. This represses global translation initiation and induces specific stress response proteins. In the RSR pathway, ZAKα senses ribosome collisions and activates p38 and JNK kinases. Therefore, we investigated the impact of suppressing each ribosome collision sensor on SINV replication. In parallel, we inactivated the *ASCC3* gene using a second sgRNA, which confirmed an inhibitory effect on SINV propagation similar to that shown in Figure 5A (Figure 6F). We found that GCN1, but not ZNF598 or ZAKα, was necessary for viral spread (Figure 6F). Consistent with this, the amount of capsid decreased in GCN1-depleted cells, while remaining unchanged in ZNF598-depleted cells (Figure 6G). Additionally, SINV infection did not induce ZAKα-mediated phosphorylation of p38, unlike UV-C treatment which was used as a positive control (Figure 6H). These findings suggest that GCN1, but not ZNF598 or ZAKα, is critical for SINV. Lastly, we investigated the importance of uS3/uS5 ubiquitin marks for the virus. These stalled ribosome marks are catalyzed by RNF10 and removed by USP10^60,61^ (Figure 6I). Suppressing or increasing them by deleting RNF10 or USP10, respectively, did not alter the number of mCherry-positive cells (Figure 6J) or capsid protein amounts (Figure 6K). As expected, infected cells depleted of RNF10 or USP10 displayed decreased or increased levels of ubiquitinated uS3/uS5, respectively (Figure 6K). By contrast, suppressing ZNF598 or ASCC3 did not affect ubiquitinated uS3 levels (Figure 6L). Collectively, our results suggest that SINV infection is linked to stalled, not collided, ribosomes. Ribosomes stall when they encounter difficult-to-translate sequences or when they are defective. Interestingly, a recent study showed that GCN1/GCN2, RNF10, and RIOK3 work together to eliminate decoding-incompetent ribosomes^65^. Such a mechanism might occur in SINV-infected cells, where we found that: 1) GCN1 is essential for SINV, 2) RNF10 ubiquitinates uS3/uS5, and 3) RIOK3 is enriched in the polysome interactome of SINV-infected cells (Figure 3C). Furthermore, we discovered SINV hijacks ASCC3. However, its role seems unrelated to its main function in resolving ribosome collisions.

### The ASCC3 helicase plays a role in the secretory pathway by promoting the synthesis of specific signal peptide-containing proteins

Next, we examined the impact of suppressing ASCC3 on the proteome of mock- and SINV-infected cells. We found that, of the ∼6,200 proteins detected, 57 proteins increased and 111 decreased significantly in uninfected ASCC3-depleted cells compared to control cells (Table S9). GO analyses revealed that proteins with a signal peptide were enriched among those decreased by ASCC3 suppression (Figure 7A). Of the 111 decreased proteins, 55 contain a signal peptide (colored turquoise in Figure 7B). A total of 291 proteins with a signal peptide were detected in the entire proteome, implying that suppressing ASCC3 affects ∼19% of signal peptide-containing proteins. This proportion even rises to 26% (76 out of 291 proteins) in SINV-infected cells (Table S9). Notably, most proteins exhibit similar variations in ASCC3-deficient cells, regardless of whether they are mock- or SINV-infected (Extended Figure 7A). The effect on signal peptide-containing proteins was confirmed using a second sgRNA for *ASCC3* gene editing (Extended Figure 7B and Table S10), but not when ZNF598 was deleted (Figure 7C, Extended Figure 7C, and Table S11). Additionally, we examined viral protein synthesis in ASCC3-knockout cells. While the levels of non-structural and mCherry proteins remained unchanged, the levels of structural proteins decreased (Figure 7D). Interestingly, the nascent polyprotein that gives rise to viral structural proteins contains a signal peptide and is translocated to the ER. This further supports that ASCC3 can be involved in the synthesis of certain signal peptide-containing proteins. GO analyses also revealed an enrichment of ER proteins and glycoproteins among those decreased by ASCC3 suppression (Figure 7A). Consistent with this finding, we observed a 20% reduction in global glycoprotein levels in ASCC3-deficient cells relative to control cells (Figure 7E). Cell fractionation experiments coupled with immunoblotting revealed that ASCC3 is localized in the cytosol, as expected, but also in the ER (Figure 7F). Control immunoblots confirm the exclusive localization of STT3A and SRPRA in the ER, and their levels decreased in ASCC3-depleted cells, which agrees with proteomic results (Figure 7F and Table S9). Fractionation experiments on SINV-infected cells revealed that viral infection did not alter the even distribution of ASCC3 between the cytosol and the ER (Figure 7G). Furthermore, stalled ribosomes were predominantly located in the cytosol rather than the ER of infected cells (Figure 7G), suggesting that the majority of them are not engaged in the cotranslational translocation of secretory and membrane proteins into the ER.

**Figure 7.**
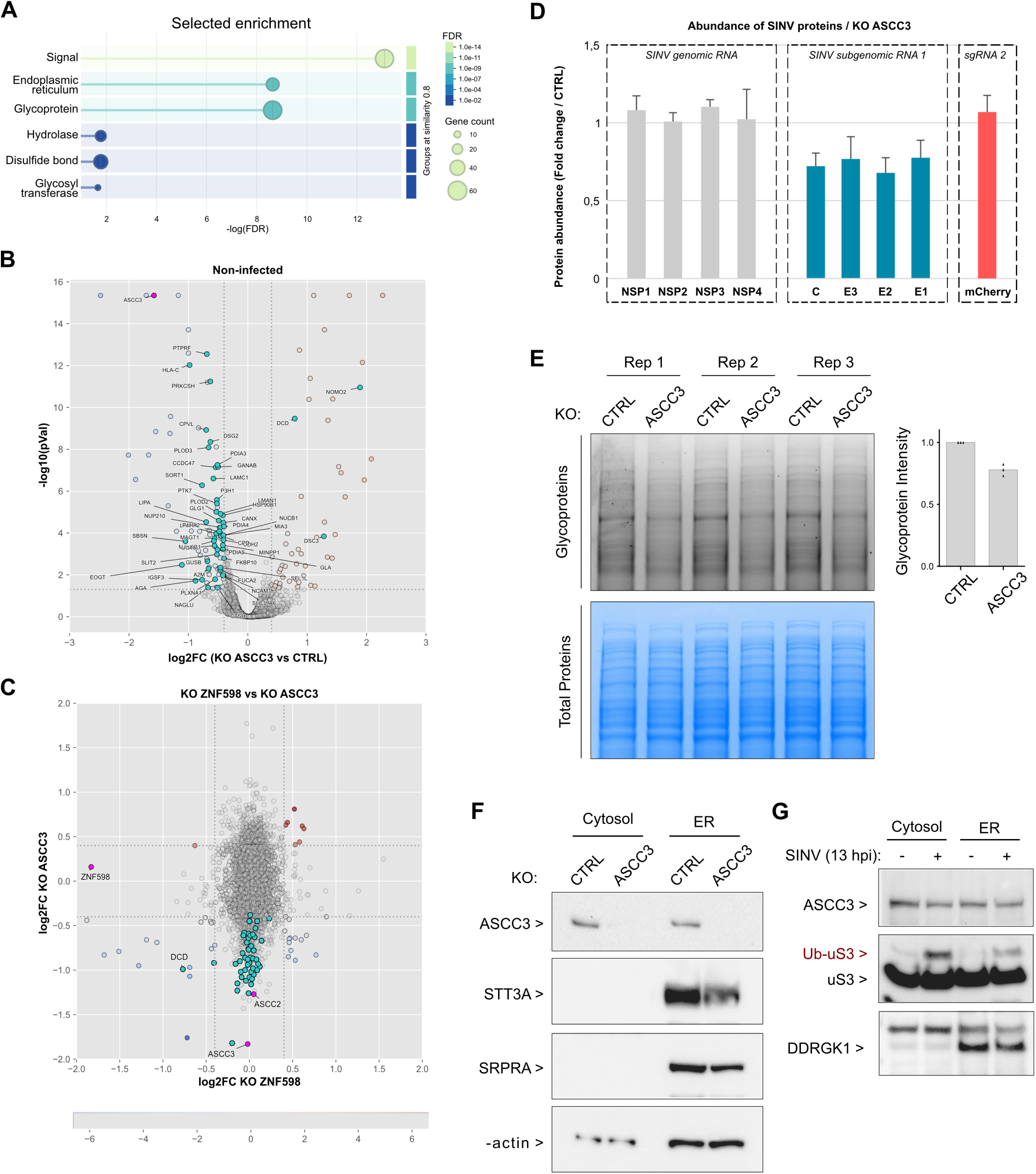
(A) Functional enrichment analysis of *ASCC3*-knockout cells using the STRING database^69^. (B) Volcano plot showing changes in protein abundance in *ASCC3*-knockout cells compared to control knockout cells. Proteins containing a signal peptide are colored turquoise and ASCC3 is colored fuchsia. (C) Scatterplot showing the relationship between changes in protein abundance in *ZNF598*-knockout cells (x-axis) and *ASCC3*-knockout cells (y-axis). Proteins containing a signal peptide are colored turquoise. ZNF598, ASCC3, and ASCC2 are colored fuchsia. (D) Quantification of all SINV proteins in cells lacking ASCC3 by mass spectrometry, compared to control cells. The cells were infected at a high MOI for 13 hours. (E) Cell lysates from control- and *ASCC3*-knockout cells were separated by polyacrylamide gel electrophoresis. The glycoproteins were stained with Pro-Q Emerald 488 dye, and the resulting green fluorescent signals were imaged (see upper panel). Total proteins were subsequently stained with Coomassie Blue (lower panel). The bar plot shows the quantification of glycoprotein signal intensity and represents the mean of three biological replicates. (F) Control and *ASCC3*-knockout cells were subjected to subcellular fractionation of the cytosol and membranes. An immunoblot was performed to analyze the subcellular localization of ASCC3. The quality of the cell fractionation was verified by immunoblotting the ER proteins STT3A and SRPRA. The β-actin immunoblot serves as a loading control. (G) Cells that were either left uninfected or infected with SINV at a high MOI for 13 hours were subjected to subcellular fractionation of the cytosol and membranes. An immunoblot was performed to analyze the subcellular localization of ASCC3 in response to viral infection. An immunoblot against ubiquitinated uS3 was performed to evaluate the partitioning of stalled ribosomes between the cytosol and the ER. The quality of the cell fractionation was verified by immunoblotting the ER protein DDRGK1.

Finally, *in situ* hybridization analyses were performed to compare the localization of ASCC3 with that of the SINV subgenomic RNA. This viral RNA encodes a precursor polyprotein that is cotranslationally translocated to the ER for processing, and it is the predominant SINV RNA at 12 hpi^26^. We observed that ASCC3 partially colocalizes with the subgenomic RNA in cytoplasmic foci (Figure 8A). These foci correspond to viral factories and are located in close proximity to the ER^26^. Importantly, suppressing ASCC3 altered the formation of these foci, producing a more diffuse viral RNA staining pattern in the cytoplasm (Figure 8A). Quantifying the fluorescent signals in a large number of cells revealed that ASCC3 and SINV RNA signals overlap within the cytoplasm (Figure 8B), finding a significant correlation coefficient (R = 0.66) (Figure 8C). These results further emphasize the importance of ASCC3 for SINV. One possibility is that ASCC3 plays a role in the synthesis and maturation of viral glycoproteins, as suggested by ASCC3’s unanticipated role in promoting the synthesis of specific proteins with signal peptides.

**Figure 8.**
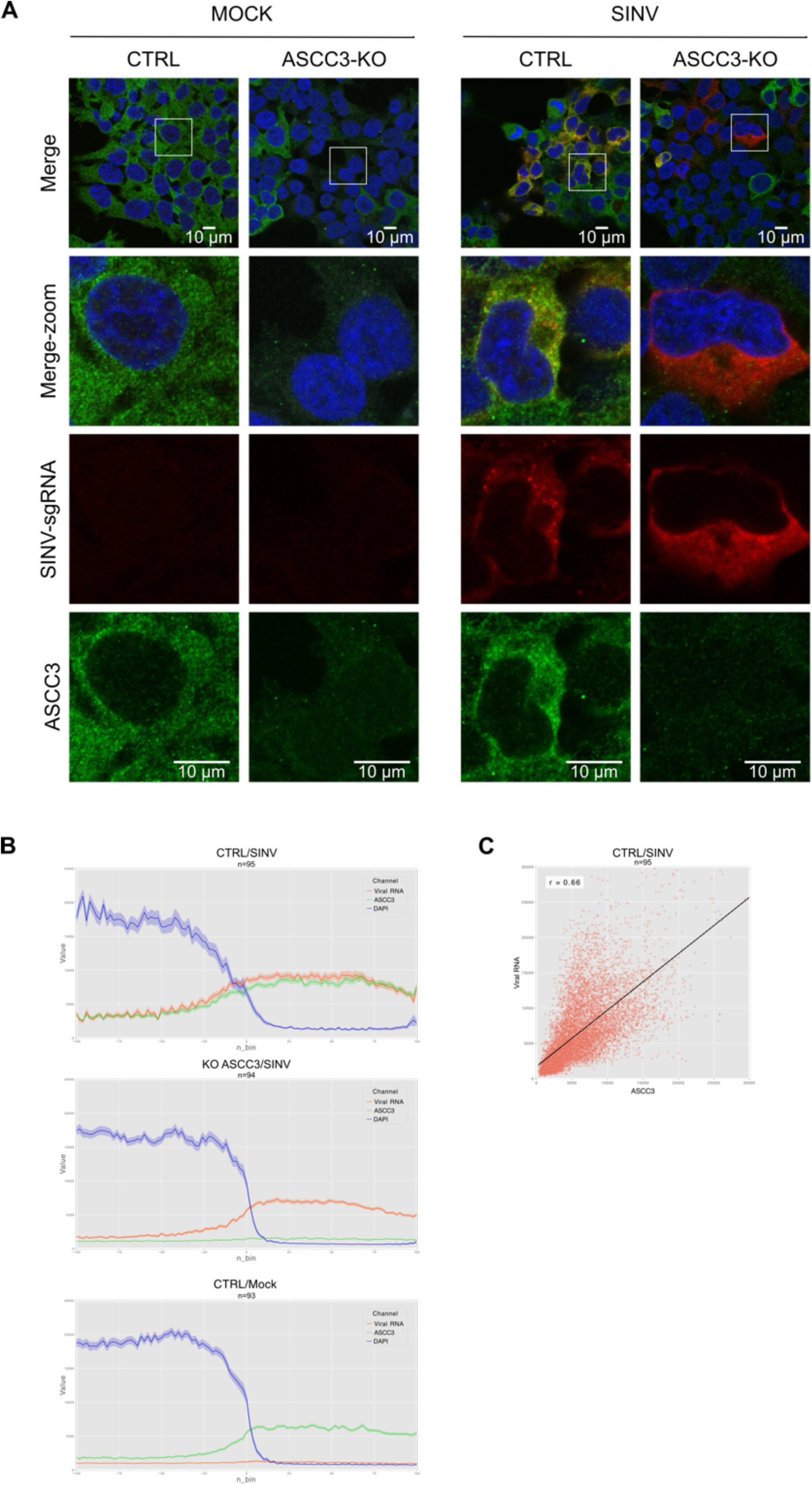
(A) Representative images showing the localization of ASCC3 and SINV subgenomic RNA by combined immunofluorescence and *in situ* hybridization in HEK293T cells. Control and *ASCC3*-knockout cells were either mock-infected or infected with SINV for 12 hours. (B) Fluorescence intensities of ASCC3, SINV RNA, and nuclear signals measured in over 90 cells shown in panel (A). (C) Correlation between ASCC3 and SINV RNA fluorescence signals in infected control cells, with a calculated Pearson coefficient of R = 0.66.

## Discussion

Our study demonstrates that eRAP is a powerful method for efficiently purifying cytoplasmic ribosomes and their associated proteins in human cells, while excluding nuclear precursors and mitochondrial ribosomes. This crosslinking-based purification approach may be valuable for discovering and analyzing ribosome-centric regulatory networks in various biological contexts. Many of the RAPs identified here do not have a known function in translation and will require further investigation to determine their role, if any, in the context of their ribosome association. Our findings further support the growing view that the translating ribosome is a dynamic regulatory hub that controls various processes rather than a passive actor in translation^66,67^. Our extensive collection of RAPs also lends support to the emerging idea that heterogeneous ribosomes can form dynamically to carry out specific functions^8^. Interestingly, many RAPs correspond to known RNA-binding proteins (RBPs). However, their recruitment to ribosomes during an infection does not correlate with changes in their binding to mRNAs. This suggests that their association with ribosomes is mostly regulated independently of their RNA-binding activity, raising the possibility that RBPs could have different functions when binding to ribosomes than when binding to mRNAs. Alternatively, their association with ribosomes may act as a decoy mechanism, temporarily sequestering them away from mRNAs to modulate post-transcriptional gene regulation during infection.

We reasoned that viruses might induce the formation of heterogeneous ribosomes optimized for translating viral RNAs and applied the eRAP method to two diverse viral model systems: HSV-1, a large DNA virus that replicates within the host cell nucleus, and SINV, a positive single-stranded RNA virus that replicates within the cytoplasm. We found that although neither HSV-1 nor SINV significantly alters the composition of the core ribosome, both viruses substantially reshape the RAP repertoire during infection. Additionally, most changes in RAP abundance are unique to the two viruses examined, supporting the notion that viruses can induce ribosome heterogeneity and specialization at the RAP level. Examining other virus model systems could provide further insight into this topic and help identify valuable targets for new antiviral therapies.

Further research on SINV revealed significant changes in the ribosome interactome that are crucial for viral propagation. Specifically, SINV promotes the association of the OST complex and PDIA6 with ribosomes, likely facilitating cotranslational N-glycosylation and folding of viral envelope proteins. SINV infection also increases DENR-MCTS1 binding to ribosomes. Conversely, SINV infection results in a marked decrease of eIF2 in the ribosome interactome. eIF2 is the primary canonical translation initiation factor that loads the initiator tRNA onto 40S ribosomes. Our findings strongly suggest that DENR-MCTS1 may replace eIF2 and drive the translation of viral subgenomic RNA, which is consistent with results of previous *in vitro* experiments^55^. We also found that SINV enhances the binding of the RQT complex to ribosomes. The core member of this complex, ASCC3, is responsible for resolving ribosome collisions through its helicase activity^32,33^. However, in the context of SINV infection, we discovered a new role for ASCC3 that appears independent of ribosome collisions and ZNF598 activity. Specifically, ASCC3, but not ZNF598, is necessary for synthesizing certain proteins with a signal peptide, including the SINV polyprotein, which gives rise to viral structural proteins. The distinct roles of ASCC3 and ZNF598 are reminiscent of a previous study showing the importance of ASCC3, but not ZNF598, in cells treated with small molecules that selectively stall translation^68^. These stalling events were reported to occur near the N-terminus of the targeted proteins^68^. Interestingly, closer examination of their amino acid sequences reveals that most contain signal peptides or membrane domains. Our data suggest that ASCC3, in addition to resolving ribosome collisions sensed by ZNF598^58,32,33^, could also be involved in resolving selective stalled ribosomes, depending on nascent chain sequence context. Further research integrating transcriptome and translatome-wide studies is necessary to define the underlying mechanism.

## Acknowledgments

We acknowledge Adeline Page from Protein Science Facility of SFR Biosciences (Université Claude Bernard Lyon 1, CNRS UAR3444, Inserm US8, ENS de Lyon) for mass spectrometry analysis. We gratefully acknowledge support from the CNRS/IN2P3 Computing Center (Lyon) for providing computing and data-processing resources needed for this work. We acknowledge the contributions of the CELPHEDIA Infrastructure (http://www.celphedia.eu/), especially the center AniRA in Lyon for flow cytometry. We gratefully acknowledge support from the PSMN (Pôle Scientifique de Modélisation Numérique) of the ENS de Lyon for all computing resources. Quantitative proteomic experiments were partially supported by Agence Nationale de la Recherche under projects ProFI (Proteomics French Infrastructure, ANR-10-INBS-08 & ANR-24-INBS-0015) and GRAL, a program from the Chemistry Biology Health (CBH) Graduate School of University Grenoble Alpes (ANR-17-EURE-0003). This work was funded by Agence Nationale des Recherches sur le SIDA et les Hépatites Virales (ANRS – ECTZ3306), Fondation FINOVI, Sidaction, the European Research Council (ERC-StG-LS6-805500) under the European Union’s Horizon 2020 research and innovation programs and the ATIP-Avenir program to E.P.R. T.J.M.S benefited from a fourth year PhD grant by Fondation pour la recherche médicale (FRM). The funders had no role in study design, data collection and interpretation, or the decision to submit the work for publication.

## Materials & Methods

### Cell culture and transfection

Baby hamster kidney cells (BHK-21) were used to grow virus stocks and were maintained in DMEM supplemented with 10% fetal bovine serum (FBS) and antibiotics, at 37 °C with 5% CO_2_. Human embryonic kidney cells (HEK293T) were maintained under the same conditions for no more than 20 passages, and were repeatedly screened for mycoplasma.

To generate the HEK293T cell line with the tagged uS7 ribosomal protein, genome editing was achieved by using the CRISPR/Cas9 technology. A single guide RNA (sgRNA) targeting the uS7 locus near the translation start codon was cloned into the PX459 expression plasmid bearing both sgRNA scaffold backbone (BB) and Cas9 nuclease. pSpCas9(BB)-2A-Puro (PX459) V2.0 was a gift from Feng Zhang (Addgene plasmid #62988; http://n2t.net/addgene:62988; RRID:Addgene_62988)^70^. Sequence of the sgRNA targeting uS7 is 5’GCCTGTCCCAGGATGACCGAG3’. Sequence of the single-stranded DNA donor template (ssODN) used for inserting the 8xHis-Flag tag sequence by homology-directed repair downstream of the uS7 translation start codon is:5’TGGGAATGAGTGCGCCTTTGCTCCATGCTAGCTGAGCTCTGACGTTTTTTTCCTGTCATCACCCTGTCCCAGGATGCACCATCACCATCACCATCACCATGACTACAAAGACGATGACGACAAGACCGAGTGGGAGACAGCAGCACCAGCGGTGGCAGAGACCCCAGACATCAAGCTCTTTGGGAAGTGGAGCACCGATG3’. The 8xHis-Flag-uS7 HEK293T cell line was obtained by transfecting the parental cells with the PX459 plasmid containing the uS7 sgRNA, as well as the aforementioned uS7 ssODN. A clonal cell population was then isolated by limiting dilution. We analyzed transgene integrity by sequencing amplicons that were subcloned into a plasmid using a TOPO TA Cloning Kit for Subcloning (Invitrogen). We verified the expression of the tagged uS7 protein by immunoblotting with an anti-Flag antibody.

To generate transient knockout HEK293T cells, the parental cells were transfected with a Cas9-expressing plasmid and a pBLADE plasmid expressing an sgRNA targeting the locus of interest near the translation start codon (See Table S11 for sgRNA sequences). The transfection was performed using jetOPTIMUS® reagent.

### Viruses

The HSV-1 strain 17 syn^+^ was provided by Dr. Anna Salvetti. HSV-1 was produced and titrated using standard procedures in BHK-21 cells. Forty-eight hours after infection, the cell supernatant was removed, the cells were collected and were frozen in liquid nitrogen. Cells were then subjected to 3 successive freeze-thaw cycles to release the viral particles. The viral suspension was aliquoted and stored at −80 °C.

The plasmid that encodes the Sindbis virus (SINV) genome with an mCherry reporter was provided by Alfredo Castello^26^. To produce the virus, the pT7-SVmCherry plasmid was linearized with XhoI and used as a template for *in vitro* RNA transcription with a HiScribe® T7 ARCA mRNA kit (New England Biolabs). The transcribed RNA was then transfected into BHK-21 cells using Lipofectamine^TM^ 3000 reagent (Invitrogen). Virus-containing supernatant was collected 24 hours after transfection, centrifuged at 1000g for 5 min, and cleared using a 0.45 µM filter. The cleared viral suspension was supplemented with glycerol (10% final), aliquoted, and stored at −80 °C. Virus titration was performed using HEK293T cells infected with serial dilutions of the viral suspension. After 24 hours, the cells were fixed with 4% paraformaldehyde, and the mCherry fluorescence was analyzed by flow cytometry using a MACSQuant VYB analyzer (Miltenyi Biotec) and quantified using FlowJo software. For most experiments, infections were performed at a MOI of 3-5 to ensure that more than 90% of the cells were infected. For functional experiments, cells were infected at a lower MOI to ensure that no more than 20-30% of the cells were infected.

### enhanced Ribosome Affinity Purification (eRAP) method

HEK293T cells engineered with 8XHis-Flag-RPS5, as well as parental (wild-type, WT) cells used as a control, were seeded into 15-cm plates at a density that made them 80% confluent at harvest. The cell culture medium was replaced with ice-cold PBS supplemented with 100µg/mL of cycloheximide and 0.1% formaldehyde (w/v) to crosslink the cells. Plates were immediately placed at 4 °C for 10 min. Crosslinking was quenched by replacing the formaldehyde solution with ice-cold PBS supplemented with 250 mM glycine at 4 °C for 10 min. Cells were scraped, collected in an ice-cold Eppendorf tube, and pelleted by centrifugation at 500 g for 5 min at 4 °C. Cells were lysed in 25 mL of lysis buffer (25 mM Tris, pH 7.4; 150 mM NaCl; 15 mM MgCl_2_; 10 mM CaCl_2;_ 1% Triton X-100; 1 mM DTT; 12 U/mL Nuclease S7 (Roche); 0.2X cOmplete^TM^ EDTA-free protease inhibitor cocktail (Roche)) for 10 min at 4 °C. The lysates were cleared by centrifugation at 1,300 g for 15 min at 4 °C. The recovered supernatants were each mixed with 300 µL of prewashed anti-Flag M2 Affinity Gel (Sigma) in washing buffer (25 mM Tris, pH7.4; 150 mM NaCl; 15 mM MgCl_2_; 1% Triton X-100) and incubated at 4 °C with gentle agitation for 1 h. Then, samples were loaded onto an Econo-Pac gravity-flow chromatography column (Bio-Rad), and the retained agarose beads were washed four times with 25 mL of washing buffer. Next, beads were transferred to 2-mL Protein LoBind tubes (Eppendorf) and washed three times with 1 mL of washing buffer. Beads were resuspended in washing buffer supplemented with 500 µg/mL of Flag peptide (DYKDDDDK peptide, GenScript), transferred to 1.5-mL Protein LoBind tubes (Eppendorf), and incubated at 4 °C with gentle agitation for 30 min. The agarose beads were removed by centrifuging the Flag eluates through a Mini Bio-Spin Chromatography Column (Bio-Rad) at 100 g and 4 °C for 1 min. For tandem affinity purification, 50 µL of prewashed Dynabeads^TM^ His-Tag Isolation and Pulldown (Invitrogen) were added to the Flag eluates and incubated at 4 °C with gentle agitation for 30 min. After discarding the supernatant, the magnetic beads were resuspended in 400 µL of washing buffer, transferred to a 2-mL Protein LoBind tube, and washed for 5 min at 4 °C. After three additional washes, the beads were transferred to a new 2-mL Protein LoBind tube. For LC-MS/MS analyses, the beads were washed twice more with washing buffer without Triton X-100. Purified proteins were eluted by adding 200 µL of His elution buffer (500 mM imidazole, pH 6.0; 500 mM NaCl) and incubating at room temperature with gentle agitation for 45 min. The eluates were incubated at 70 °C for 15 min to reverse the formaldehyde crosslinking and stored at −20 °C prior to LC-MS/MS analysis or immunoblotting.

### Polysome fractionation on sucrose density gradient centrifugation

For polysome fractionation, the cells were seeded the day before and harvested at 80% confluence. After removing the culture medium, cells were washed with ice-cold PBS supplemented with 100 µg/mL cycloheximide. If the cells were crosslinked before lysis, the culture medium was replaced with ice-cold PBS containing 0.1% formaldehyde (w/v) and 100 µg/mL cycloheximide and the plates were placed at 4 °C for 10 min. The supernatants were removed and the crosslink was quenched by adding ice-cold PBS supplemented with 250 mM glycine and 100 µg/mL cycloheximide at 4 °C for 10 min. Cells were scraped, collected in 1.5-mL Eppendorf tube, and centrifuged at 500g and 4 °C for 5 min. Cells were lysed in 1 mL of lysis buffer (25 mM Tris, pH 7.4; 150 mM NaCl; 15 mM MgCl_2_; 1 mM DTT; 100 µg/mL cycloheximide; 1% Triton X-100; 1X cOmplete^TM^ EDTA-free protease inhibitor cocktail). Lysates were homogenized by gently pipetting up and down eight times with a P1000 pipettor and then incubated at 4 °C for 10 min. Lysates were cleared by centrifugation at 1,300g and 4 °C for 10 min. The resulting supernatants were layered on top of 10-50% (w/v) linear sucrose gradients prepared in 20 mM HEPES-KOH, pH 7.4, 5 mM MgCl_2_, 100 mM KCl, 2 mM DTT, and 100 µg/mL cycloheximide. The Gradient Master (Biocomp) was used for making the gradients. Gradients were centrifugated in a SW 41Ti rotor at 35,000 rpm for 2 h 40 min at 4 °C and fractionated from the top of the gradient with continuous monitoring of the absorbance at 254 nm. The fractions were analyzed by immunoblotting or tandem affinity purification of polysomes.

### eRAP method applied to pooled polysomes

For tandem affinity purification of polysomes, the polysomes were fractionated as described in the previous section. The polysomal fractions corresponding to disomes and higher were pooled, and the volume was adjusted to 25 mL with lysis buffer (25 mM Tris, pH 7.4; 150 mM NaCl; 15 mM MgCl_2_; 10 mM CaCl_2;_ 1% Triton X-100; 1 mM DTT; 12 U/mL Nuclease S7; 0.2X cOmplete^TM^ EDTA-free protease inhibitor cocktail). Tandem affinity purification was performed as described in the eRAP section above.

### Measurement of protein synthesis using O-propargyl-puromycin (OPP)

Cells were incubated with 10 µM OPP (Immagina Biotechnology) at 37 °C for 30 min and then fixed in 4% paraformaldehyde in PBS at room temperature for 20 min. After a permeabilization in PBS-0.5% Triton X-100 for 20 min, the cells were labelled with Click-iT^TM^ Plus Alexa Fluor^TM^ 488 Picolyl Azide Toolkit (ThermoFisher Scientific) according to the manufacturer’s instructions. The cells were then analyzed using a MACSQuant VYB cytometer (Miltenyi Biotec), and the results were quantified with FlowJo software.

### Preparation of total lysates and subcellular fractions

Total cell lysates were typically prepared using RIPA buffer (50 mM Tris, pH 7.4; 150 mM NaCl; 1% IGEPAL; 0.5% Na deoxycholate; 0.1% SDS) supplemented with 1X cOmplete^TM^ EDTA-free protease inhibitor cocktail (Roche). Cells were incubated in RIPA buffer at 4 °C for 15 min, and lysates were cleared by centrifugation at 18,000g and 4 °C for 10 min. The resulting supernatants were collected, frozen on dry ice, and stored at −80 °C prior to processing.

Subcellular fractionation of the cytosol and membranes was based on sequential detergent extraction, as previously described^71^. All steps were carried out on ice and with ice-cold reagents. In brief, cells seeded in a 10-cm plate were washed with 5 mL of PBS containing 100 µg/mL of cycloheximide, and then incubated for 5 min with 1 mL of permeabilization buffer (110 mM KOAc; 20 mM HEPES-KOH, pH 7.2; 5 mM MgCl_2_; 1 mM EGTA; 0.03% digitonin; 1 mM DTT; 100 µg/mL cycloheximide). The supernatant containing the cytosol was collected. Cells were washed with 2.5 mL of washing buffer (same as permeabilization buffer but with 0.002% digitonin) and incubated for 5 min with 1 mL of lysis buffer (200 mM KOAc; 20 mM HEPES-KOH, pH 7.2; 5 mM MgCl_2_; 1 mM EGTA; 2% n-Dodecyl β-D-maltoside; 1 mM DTT; 100 µg/mL cycloheximide). The supernatant containing the membrane fraction was collected. The cytosolic and membrane fractions were both cleared at 12,000g and 4 °C for 5 min to remove cell debris. The resulting supernatants were recovered, frozen on dry ice, and stored at −80 °C prior to processing.

### Protein analysis

Silver staining of protein gels was performed using a Pierce^TM^ silver stain kit (ThermoFisher Scientific). Protein concentrations were measured using a Pierce^TM^ BCA Protein assay kit (ThermoFisher Scientific). For immunoblot analysis, 30 µg of proteins were typically mixed with 1X Bolt LDS sample buffer and 1X Bolt sample reducing agent (both from Invitrogen). The samples were denatured at 70 °C for 10 min, and loaded onto Bolt Bis-Tris Plus gels (Invitrogen). After electrophoresis in Bolt MES buffer (Invitrogen), proteins were transferred to a Hybond P 0.45 PVDF membrane (Cytiva). Membranes were blocked for 1 h at room temperature in 5% nonfat dry milk in TBS-Tween (0.1%Tween 20), and then incubated overnight at 4 °C with primary antibodies diluted in TBS-Tween containing 3% bovine serum albumin (BSA). All antibodies were diluted according to the manufacturer’s instructions and are listed in the Key Resources table. Membranes were washed three times for 10 min with TBS-Tween and then incubated for 45 min with horseradish peroxidase-conjugated secondary antibody diluted in TBS-Tween. After three 10-min washes with TBS-Tween, the membranes were developed using SuperSignal^TM^ West Pico PLUS chemiluminescent substrate (ThermoFisher Scientific) and imaged with a ChemiDoc Touch imaging system (Bio-Rad). The staining of glycoproteins directly in gels was done using the Pro-Q^TM^ Emerald 488 glycoprotein gel and blot stain kit (Invitrogen), following the instructions provided by the manufacturer. The stained gels emitting a green fluorescent signal were imaged with a ChemiDoc MP imaging system (Bio-Rad).

### Mass spectrometry analyses

#### Ribosome interactomes

Proteins from ribosomal preparations obtained from various formaldehyde percentages were solubilized in Laemmli buffer before being stacked in the top of a 4–12% NuPAGE gel (Life Technologies) stained with R-250 Coomassie blue (Bio-Rad) and in-gel digested using modified trypsin (sequencing grade, Promega) as previously described^72^. Proteins from ribosomal preparations obtained during infection with HSV-1 or SINV were digested using SP3 magnetic beads^73^ and desalted using Ultramicro spincolumn C18 (protocol from manufacturer Harvard Apparatus). The dried extracted peptides were resuspended in 5% acetonitrile and 0.1% trifluoroacetic acid and analyzed by online nanoliquid chromatography coupled to tandem mass spectrometry (LC–MS/MS) (UltiMate 3000, RSLCnano and Q-Exactive HF, ThermoFisher Scientific). Peptides were sampled on a 300 μm X 5 mm PepMap C18 precolumn (ThermoFisher Scientific) and separated either on a 200 cm µPAC column (PharmaFluidics) or a 75 μm X 25 cm C18 column (Reprosil-Pur 120 C18-AQ, 1.9 μm, Dr. Maisch HPLC GmbH). The nano-LC method consisted either of a 120 min or 60 min multi-linear gradient at a flow rate of 300 nL/min, ranging from 5 to 41% acetonitrile in 0.1% formic acid. Survey full-scan MS spectra (*m*/*z* = 400–1600) were acquired with a resolution of 60,000 after the accumulation of 10^6^ ions (maximum filling time 200 ms). The 20 most intense ions were fragmented by higher-energy collisional dissociation after the accumulation of 10^5^ ions (maximum filling time: 50 ms) either with normalized collision energy of 30 or 27. MS and MS/MS data were acquired using the software Xcalibur (ThermoFisher Scientific).

Data were processed automatically using Mascot Distiller software (version 2.7.1.0, Matrix Science). Peptides and proteins were identified using Mascot (version 2.8) through concomitant searches against UniProt reference proteomes (Homo sapiens taxonomy, downloaded in July 2025 as well as HSV-1 strain 17 or SINV taxonomy), the His-Flag-uS7 protein sequence, and a homemade database for classical contaminants. Trypsin/P was chosen as the enzyme, and two missed cleavages were allowed. Precursor and fragment mass error tolerance were set to 10 ppm and 20 ppm, respectively. Peptide modifications allowed during the search were: carbamidomethylation (fixed), acetyl (protein N-terminal, variable) and oxidation (variable). The three different projects were then processed separately. Proline software (version 2.3.1)^74^ was used to merge data dependent acquisition (DDA) results. After combination, results were filtered: conservation of rank 1 peptide-spectrum match (PSM) with a minimal length of 6. Peptide-spectrum matching (PSM) score filtering is then optimized to reach a false discovery rate (FDR) of PSM identification below 1% by employing the Benjamini-Hochberg approach^75^. A minimum of one specific peptide per identified protein group was set. Proline was then used to perform MS1-based label free quantification of the peptides and protein groups from the different samples with cross-assignment. Protein abundances were computed from razor and specific peptides. The mass spectrometry proteomics data have been deposited to the ProteomeXchange Consortium via the PRIDE^76^ partner repository. Statistical analysis was performed using ProStaR 1.38.0^77^ to determine differentially abundant proteins. Protein sets were filtered out if they were not identified in at least two replicates of one condition. Protein sets were then filtered out if they were not quantified across all replicates in at least one condition. Contaminants were also filtered out. After log2 transformation and normalization (see below for parameters), partially observed values (POV) were imputed with the SLSA method and values missing in the entire condition (MEC) were imputed with 1-percentile value of each sample.

##### Ribosome interactome with various concentrations of formaldehyde

The abundance of quantified proteins was normalized using the total abundance of ribosomal proteins for each sample. An ANOVA was conducted with a p-value cut-off allowing to reach an FDR inferior to 1% according to the Benjamini–Hochberg procedure. To identify the differences between the various concentrations of formaldehyde, Prostar was used to run LIMMA test with every possible One versus One contrast. All data were merged and LIMMA p-values corrected with Benjamini–Hochberg procedure. Proteins were considered as significantly enriched (or depleted) in a condition if their FDR-adjusted p-value was below 0.01 and their log2(FC)□>□1 (or log2(FC)□<□-1). Proteins that were imputed in the up-regulated condition were manually invalidated.

#### Ribosome interactome of HSV-1-infected cells and polysome interactome of SINV-infected cells

The interactomes were initially defined by comparing samples from His-Flag-uS7 cells and from WT cells. The 0.5 quantile method was used for normalization within conditions. Prostar was used to run LIMMA test. Proteins were considered part of the interactomes if their p-value was below 0.01 and their FC > 5. For samples from His-Flag-uS7 cells, the abundance of quantified proteins (only validated in one of the interactomes) was normalized using total ribosomal protein abundance. Prostar was used to run LIMMA test. Proteins were considered as significantly enriched (or depleted) in the infected condition if their FDR-adjusted p-value was below 0.05 and their log2(FC)□>□1 (or log2(FC)□<□-1). Proteins that were imputed in the up-regulated condition were manually invalidated.

##### Ribosome interactome of SINV-infected cells

Proteins from ribosomal preparations were processed using the SP3 protocol^78^ with modifications for mass spectrometric analysis. Beads (10 μg/μL) were washed and reconstituted in 200 μL of water, and the bead mix was stored at 4°C. SP3 was performed in 1.5-mL Protein LoBind® tubes (Eppendorf). Samples were denatured in a solution of 8 M urea and 100 mM ammonium bicarbonate at pH 8, followed by reduction and alkylation with 10 mM tris-2(-carboxyethyl)-phosphine (TCEP) and 50 mM chloroacetamide. Washed beads were added to the alkylated lysates (100 μg per sample) before acetonitrile was added to a final concentration of 80%. Samples were incubated at room temperature for 30 min to allow complete binding of proteins onto beads. Subsequently, beads were immobilized using a magnetic rack for 2 min. The supernatant was discarded and beads were washed on the rack twice with 1 mL of 70% ethanol, followed by 100% acetonitrile. This washing cycle was repeated seven times. Samples were digested with trypsin (at a ratio of 1:40 [w/w] in 50 μL of 1 M urea; 50 mM triethylammonium bicarbonate; 1 mM CaCl_2_) overnight at 37°C. The digested samples were transferred to new tubes, and the beads were rinsed with 25 μL of 2% DMSO for 10 min to elute the bound peptides. The rinse was then pooled with the digested samples. Before LC-MS analysis, peptides were acidified by adding formic acid to a final concentration of 1%. Tryptic digests were subsequently analyzed on a nanoUHPLC (ThermoFisher Scientific) connected to a Q Exactive mass spectrometer (ThermoFischer Scientific) through an EASY-Spray nano-electrospray ion source (ThermoFischer Scientific). The peptides were trapped on a C18 PepMap100 pre-column (300 µm I.D. x 5 mm, 100 Å, ThermoFisher Scientific) using solvent A (0.1% formic acid in water). Trapped peptides were separated on in-house constructed analytical column (75 µm I.D. × 500 mm, Reprosil C18, 1.9 µm, 100 Å) using a linear gradient (length: 60 min, 15 % to 35 % solvent B composed of 0.1% formic acid and 5 % DMSO in acetonitrile, flow rate: 200 nL/min). The separated peptides were electrosprayed directly into the mass spectrometer operating in a data-dependent mode. Full scan MS spectra were acquired in the Orbitrap (scan range 350-1500 m/z, resolution 70,000, AGC target 3e6, maximum injection time 50 ms). After the MS scans, the 10 most intense peaks were selected for HCD fragmentation at 30 % of normalized collision energy. HCD spectra were also acquired in the Orbitrap (resolution 17,500, AGC target 5e4, maximum injection time 120 ms).

Protein identification and quantification were performed using Andromeda search engine implemented in MaxQuant (1.6.3.4). Peptides were searched against reference Uniport datasets: Homo sapiens (Uniprot_id: UP000005640, downloaded Nov 2016), and a custom database including all the known SINV polypeptides and a list of common contaminants provided by the software. Search settings were kept at default parameters with “match between run” method switched on. FDR was set at 1% for both peptide and protein identification. Protein intensities from MaxQuant search result were imported in RStudio (R Project) for further processing. Proteins that has less than 3 valid intensity measurements in at least one condition among 5 replicates were removed prior to downstream analysis. Intensities were log-2 transformed and normalized using variance stabilizing normalization method^79^. Samples were divided into contrast pairs that compare each SINV infection condition with non-infected condition, and statistical tests were conducted in each pair independently. For proteins within each contrast pair, if the protein was absent in one condition, missing values in that condition were imputed with minimal deterministic value method using 1% quantile of global intensities. Batch effects were identified and assessed using principle component analysis. Linear modeling and a Bayesian model-based moderated t-test were performed using the limma package^80^. The replicate number was incorporated into the model as a covariate to account for the batch effects. P-values obtained from the moderated t-statistics were adjusted using the Benjamini-Hochburg method to account for the effect of the large dataset.

#### Total proteomes

For proteomics analyses of total lysates, cells were lysed in RIPA buffer, as described in the corresponding section. For in-solution enzymatic digestion, 50 μg of the protein samples were mixed with the lysis buffer provided in the easyPep Mini Kit (#A40006, Thermo Scientific). Samples were reduced and alkylated by incubating them at 95 °C with continuous shaking at 1000 rpm for 10 min. Then, samples were digested at 37 °C for 3 h and processed according to the manufacturer’s protocol. After the cleanup step, the resulting peptides were dried, resuspended in 0.1% formic acid, and quantified using the Quantitative Fluorometric Peptide Assay (#23290, Thermo Scientific). Samples were analyzed using an Ultimate 3000 RSLCnano system (ThermoFisher Scientific) coupled to a Q Exactive HF mass spectrometer (ThermoFisher Scientific) via a nanoelectrospray ionization source. Four hundred nanograms of each peptide mixture were loaded onto a PepMap Neo C18 trap column (300 μm I.D. × 5 mm, 5 μm; Thermo Scientific) at a flow rate of 20 μL/min with a mobile phase of 2% acetonitrile, 0.05% trifluoroacetic acid in water for 3 min. The peptides were subsequently separated on an Acclaim Pepmap C18 100 nano-column (75 μm I.D. x 500 mm, 3 μm, 100 AL; Thermo Scientific), with a 100-minute linear gradient from 3.2% to 20% buffer B (A: 0.1% formic acid in water, B: 0.1% formic acid in acetonitrile), from 20% to 32% of B in 20 min, from 32% to 90% of B in 2 min, hold for 10 min, then return to initial conditions in 2 min for a total of 13 min. The total duration was set to 150 min at a flow rate of 300 nL/min. The oven temperature was kept constant at 40 °C. Samples were analyzed using a TOP15 HCD method. MS data were acquired in a data-dependent strategy that selected fragmentation events based on the 15 most abundant precursor ions in the survey scan (350-1650 Th). The survey scan resolution was 120,000 at m/z 200 Th. The ion target value for survey scans in the Orbitrap and MS^2^ modes were set to 3e6 and 1e5, respectively, and the maximum injection time was set to 60 ms for both scan modes. The HCD MS/MS spectra acquisition parameters were as follows: collision energy = 27; isolation width of 1.4 m/z; exclusion of precursors with an unknown or 1 charge state. The peptides selected for MS/MS acquisition were placed in an exclusion list for 20 s using the dynamic exclusion mode to prevent duplicate spectra. The spray voltage was set to 1800 V for positive ionization and the ion transfer tube was maintained at a temperature of 250 °C. For data analysis, proteins were identified by database searching using Sequest HT with Proteome Discoverer 2.5 and 3.2 software (Thermo Scientific) against the Homo sapiens Swiss-Prot database (release 2023-06, 20,348 sequences), the SINV protein sequences (10 sequences), the mCherry protein sequence, and a database of contaminants. The precursor and fragment mass tolerances were set to 10 ppm and 0.02 Da, respectively, and up to two missed cleavages were permitted. Oxidation (M) and acetylation (protein N-terminus) were set as variable modifications and carbamidomethylation (C) as a fixed modification. Full trypsin was selected as the digestion enzyme parameter. Peptide and protein validation were performed using Percolator, with an FDR of 1% for both. Protein quantification was done using the label-free quantification (LFQ) approach. LFQ abundance values were obtained and normalized to the total peptide amount. Protein quantitation was performed using the precursor ions quantifier node in Proteome Discoverer 2.5 and 3.2 software, with quantitation based on pairwise ratios and a t-test hypothesis. Proteins were considered to be differentially expressed between the two conditions when the fold change was greater than 2 or less than 0.5 and the Adj p-value was less than 0.05. The mass spectrometry proteomics data have been deposited to the Center for Computational Mass Spectrometry repository (University of California, San Diego) via the MassIVE tool.

### Gene Ontology Analysis Across Formaldehyde Concentrations

Proteins specifically pulled down using the eRAP method were first identified in the absence of formaldehyde, using cell lysates with untagged ribosomes as a control. For each formaldehyde concentration (0.025%, 0.1%, 0.37%, and 1%), a list of enriched proteins was generated relative to the condition without formaldehyde. These five protein lists were independently analyzed using StringDB to retrieve Gene Ontology (GO) annotations across the Cellular Component, Biological Process, and Molecular Function categories. The most relevant and non-redundant GO terms were manually selected. The number of proteins associated with each selected GO term was then quantified as a function of formaldehyde concentration and visualized in Figure 1E.

### Protein Clustering Across Formaldehyde Concentrations

To perform trajectory clustering, all iBAQ values from experimental replicates were considered for every protein enriched in at least one formaldehyde condition (0–1%). Each protein trajectory therefore consisted of 15 experimental values. Sixteen clusters were generated using the K-means algorithm implemented in the scikit-learn Python package. The mean trajectories of the 16 clusters are shown in Extended Figure 1A, while representative trajectories of selected protein groups are displayed in Figure 1F.

### Ribosome interactome comparison

To compare the results of our ribosome interactome obtained using the eRAP method with His-Flag-uS7 engineered cells with those obtained using the eS17-Flag-tagged cells^5^ or immunoprecipitation of rRNA^19^, we combined all enriched proteins from the three methods into a single dataset. This collection enabled the identification of proteins commonly enriched by two or more approaches, as well as those uniquely captured by each protocol. A Venn diagram was generated using the matplotlib-venn Python package to illustrate these overlaps (Figure 1G). In addition, the combined protein dataset was used to query StringDB for Gene Ontology (GO) annotations across the broadest possible scope. The number of proteins associated with each of the most relevant GO terms—particularly those related to translation—was quantified for each protocol and visualized in Figure 1H.

### Immunofluorescence and RNA FISH assays

smFISH experiments were performed as previously described^81^ with few modifications. Briefly, 2□×□10^5^ CTRL or ASCC3-KO HEK293T cells were seeded on High Precision Coverslips (Marienfeld, #0117520) in 10% FBS-supplemented DMEM for 24□h at 37□°C. Cells were either mock-infected or SINV-infected with 1 MOI in se-rum-free DMEM for 1 h at 37□°C and media was subsequently replaced with 5% FBS-supplemented DMEM. At 12□h post-infection, cells were washed with PBS and fixed in 4% formaldehyde for 15□min at room temperature. Cells were washed three times for 5□min with PBS and permeabilized with PBS supplemented with 0.2% Triton X-100 for 10□min at room temperature. Cells were washed twice with PBS for 5□min each, twice with 2x SSC for 5□min each, and twice with pre-hybridization buffer (pre-warmed, 2x SSC, and 10% deionized formamide in DEPC-treated water) for 20□min each at 37□°C. Cells were then incubated with 125□nM SINV sgRNA Stellaris probes (LGC Biosearch Technologies) in hybridization buffer (pre-warmed, 2x SSC, 10% deionized formamide, and 10% dextran sulphate in DEPC-treated water) at 37□°C overnight in a humidified chamber. From this point, cells were kept in the dark. Cells were then washed twice with pre-warmed pre-hybridization buffer for 20□min at 37□°C, twice with 2X SSC for 5□min each, and twice with 1X PBS 5□min each. Cells were incubated with 1X PBS-Tween20-0.3% (PBS-T) supplemented with BSA 5% for 30□min followed by incubation with ASCC3 primary antibody (Protein-tech 85130-2-RR 1:100 dilution) in same buffer for 1□h. Next, the cells were washed three times with PBS-T for 5□min each and incubated with secondary antibody (α-rabbit Alexa488 at 1:500 dilution) in PBS-T supplemented with 5% BSA for 1□h. The cells were washed three times in PBS-T and once in 1X PBS. Cells were incubated with DAPI (1:1000 of 1□mg/mL stock) in 1X PBS for 15□min at room temperature, followed by two washes with 1x PBS for 5□min and one wash with DEPC-treated pure water for 1□min. Finally, coverslips were mounted on glass slides using Pro-Long™ Diamond antifade mountant (ThermoFisher Scientific). Cells were imaged on a Zeiss LSM880 confocal system using a 63X oil objective (Plan-Apochromat 63x/1.4 Oil DIC M27). All solutions were RNase free and supplemented with RiboLock RNase Inhibitor (Thermo Fisher Scientific).

Acquired images were analyzed sequentially using a custom ImageJ macro followed by a Python script. Due to the high cell density required for the experimental setup, cell segmentation was not feasible. To address this limitation, fluorescence intensities for ASCC3, SINV subgenomic RNA, and nuclei were measured along a linear region of interest (ROI) drawn across the widest portion of each cell. By convention, the largest cytoplasmic region corresponded to the end of the ROI. The ImageJ macro automated the measurement of fluorescence intensity for each channel and generated corresponding CSV files. The subsequent Python script identified, for each ROI, the position marking the end of the nucleus (determined by the minimum slope of the DAPI signal). This position was then used to convert distances from pixels to percentages, with 100% representing the cytoplasmic region beyond the nucleus, enabling comparison across cells with varying geometries. Finally, the fluorescence intensity profiles of each channel were normalized and plotted from −100% to 100% (0% corresponding to the nuclear boundary) for each experimental condition (Figure 8B). The correlation between ASCC3 and SINV subgenomic RNA fluorescence signals was calculated using the pearsonr function from the scipy Python package (Figure 8C).

### Statistical information

All statistical information is detailed throughout the Figure legends and the Methods section.

## KEY RESOURCES TABLE

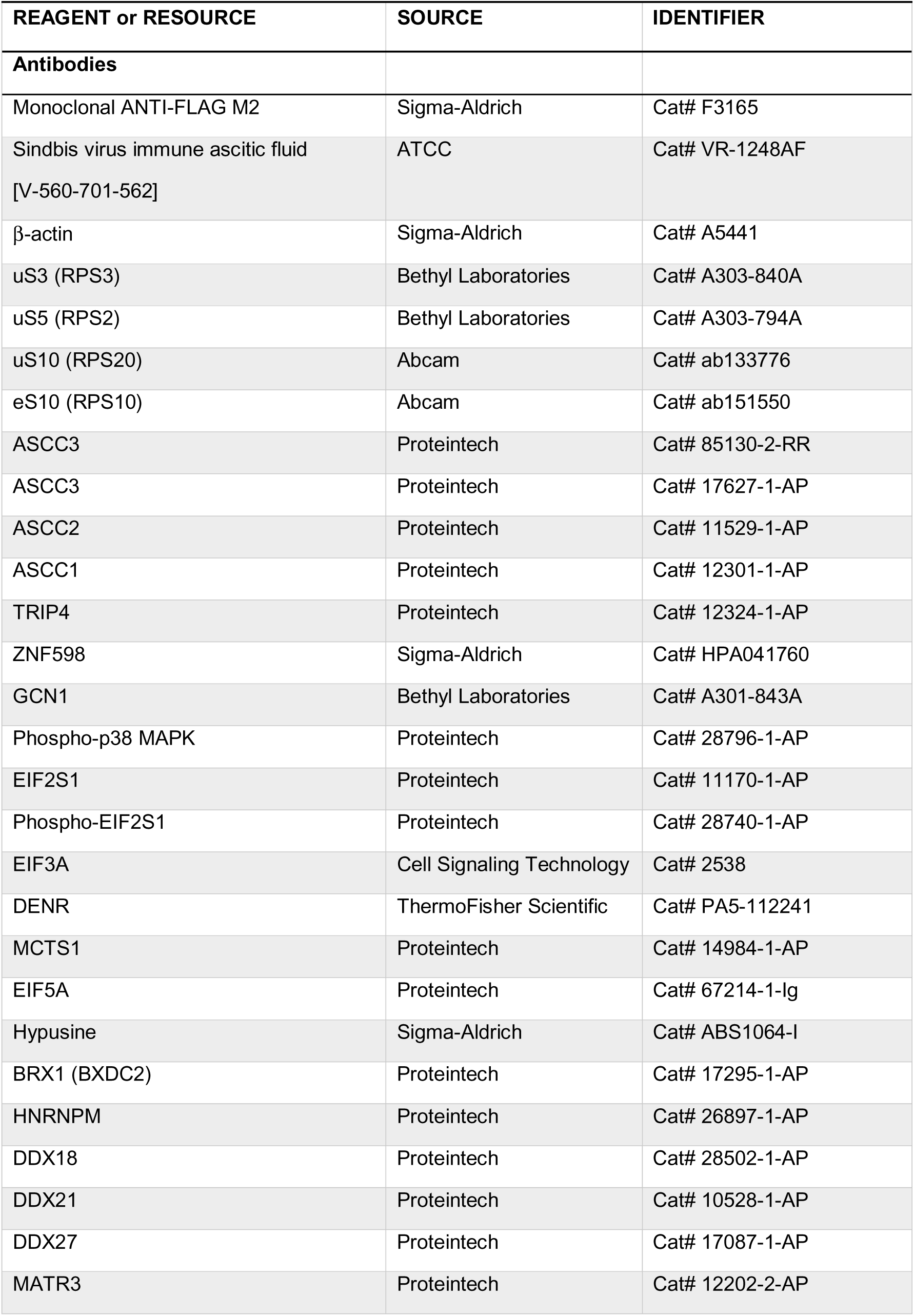
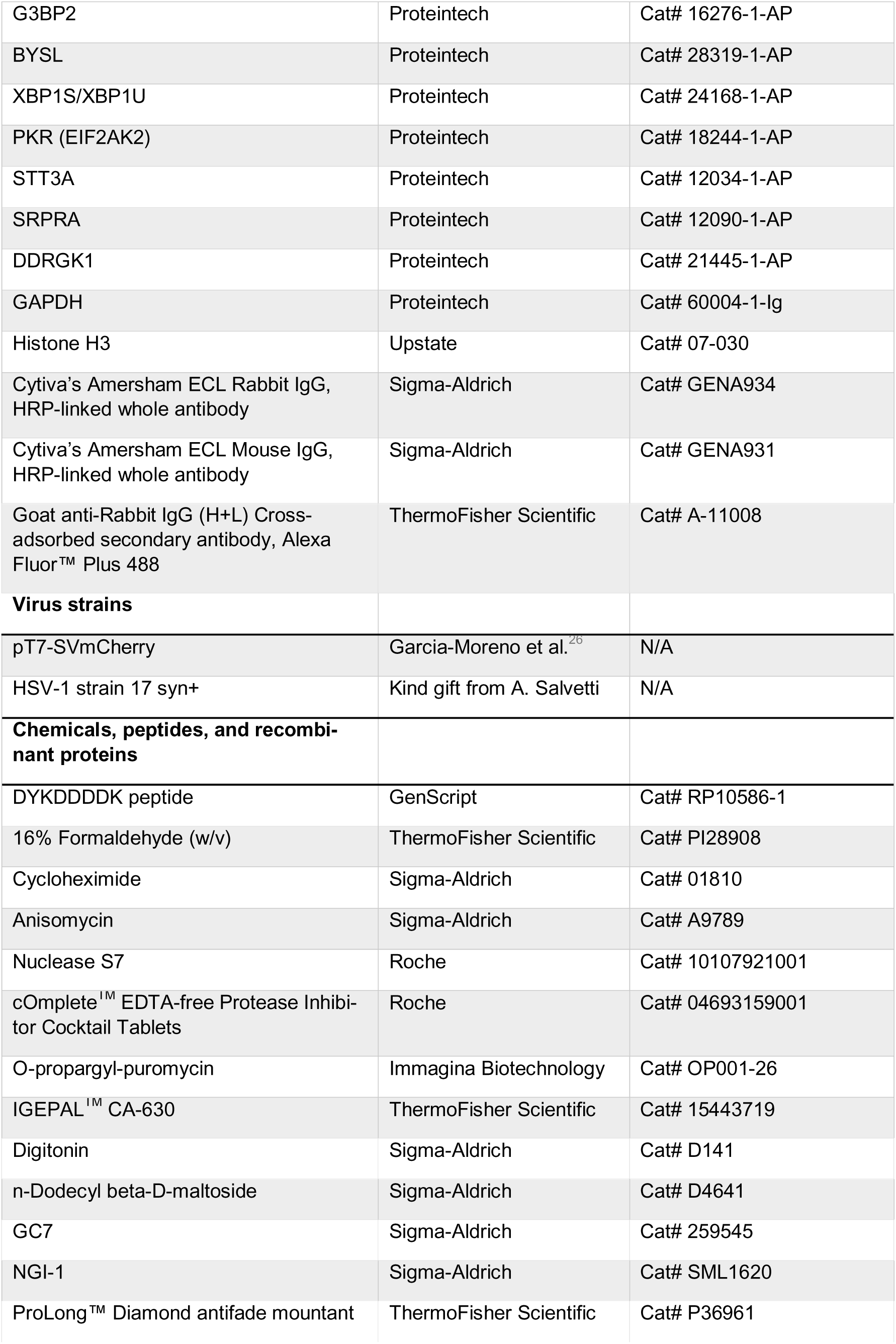
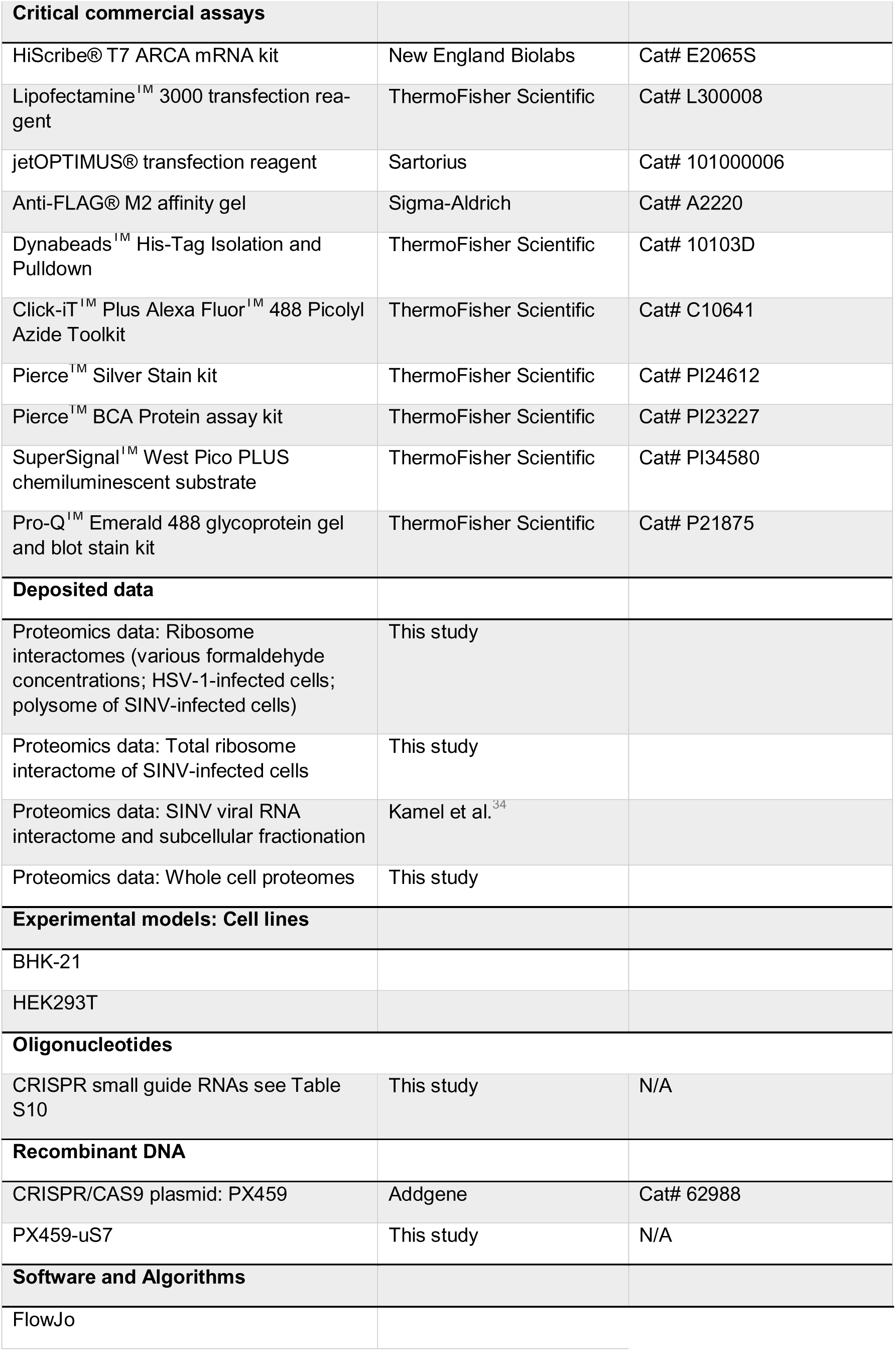

