## Supplementary figures and images for "Viral Infections Drive Functional Remodeling of Ribosome-Associated Proteins"

### Extended Figure 1

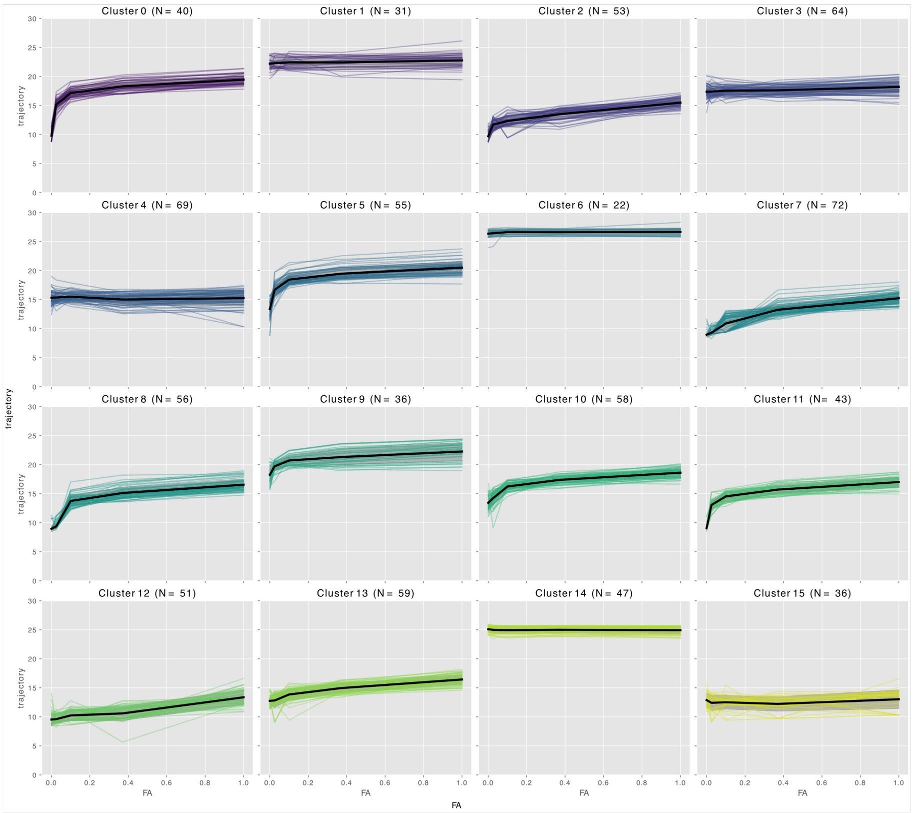

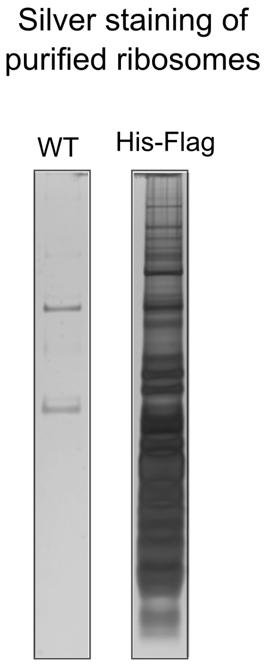

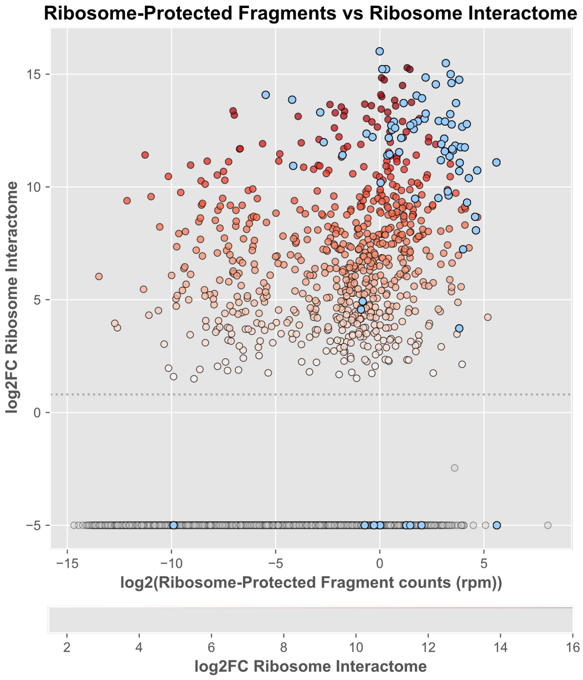

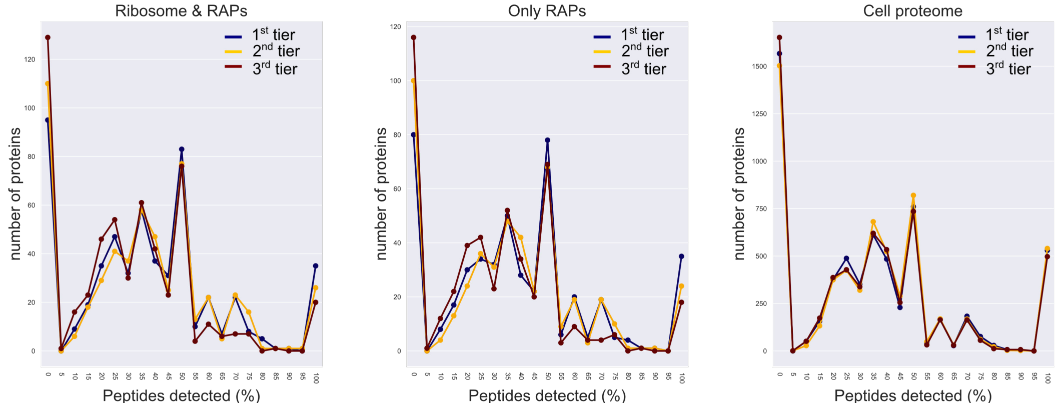

### Extended Figure 5

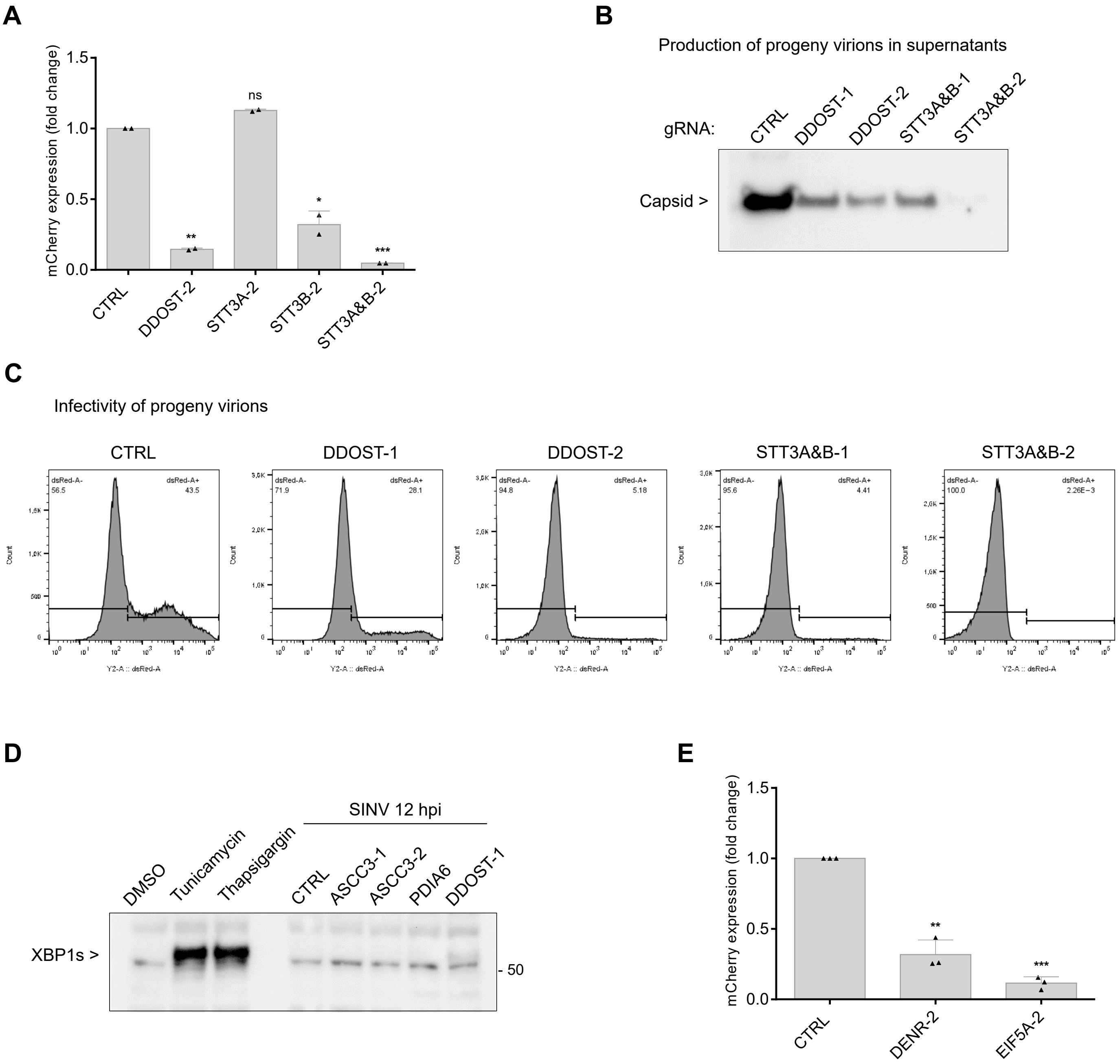

### Extended Figure 6

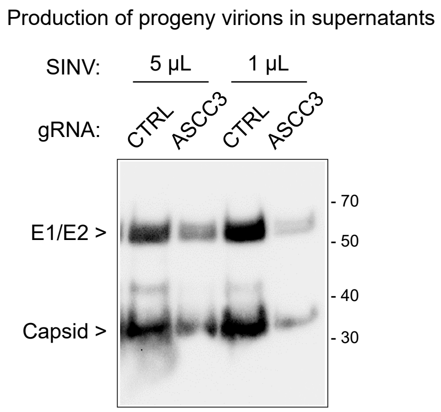

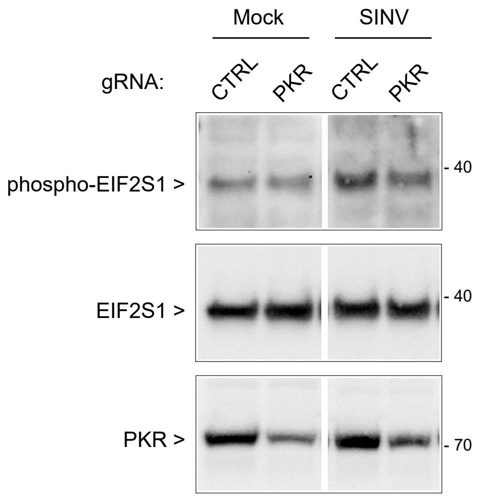

### Extended Figure 7

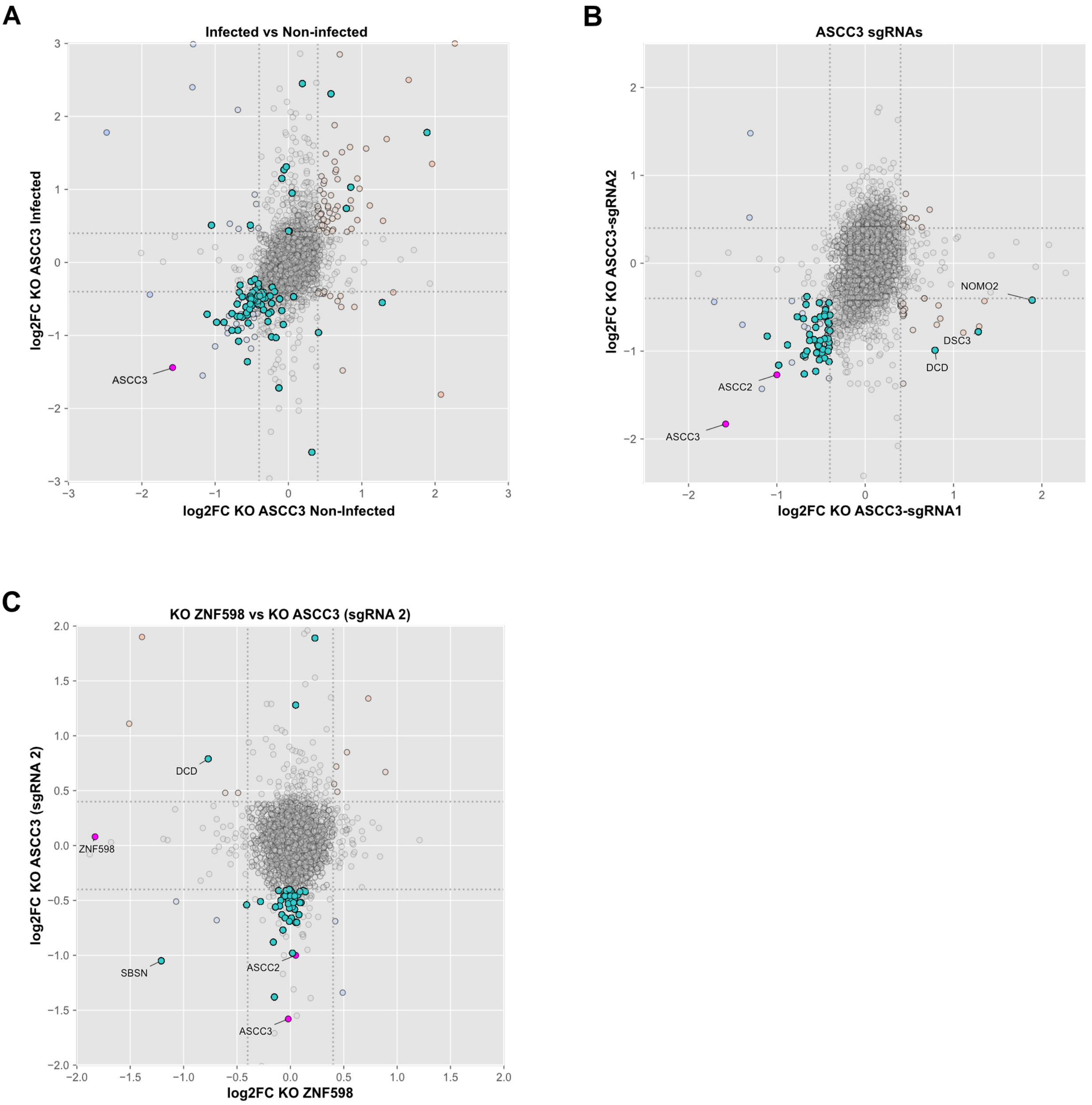
