## Supplementary Figure Legends for "Viral Infections Drive Functional Remodeling of Ribosome-Associated Proteins"

**Extended data Figure 1.**

(A) Protein clustering analysis. The black line shows the mean trajectory and the colored surface shows the standard deviation for each cluster. The N values represent the number of proteins in each cluster.

(B) Silver staining analysis of eluates from a representative eRAP experiment using HEK293T cells that express His-Flag-uS7 compared with wild-type cells.

(C) Scatter plot showing the ribosome interactome from His-Flag-uS7 cells (expressed as log_2_ fold-change relative to parental cells) and ribosome-protected fragment (RPF) counts (expressed as log_2_ reads per million). Ribosomal proteins are colored blue.

(D) Analysis of peptide distribution across the entire coding sequence of ribosomal proteins and ribosome-associated proteins. The same analysis was performed on whole cell proteome for comparison. The protein sequences were divided into tiers, and the proportion of peptides in each tier was counted. No bias toward the N-terminus was found in the peptide distribution of the ribosome-associated proteins.

**Extended data Figure 5.**

(A) Measurement of mCherry fluorescence serves as an indicator of SINV replication in HEK293T cells that were transfected with the indicated sgRNAs to edit genes encoding some OST components. The cells were infected at a low MOI for 24 hours. The bar plot shows the mean of two biological replicates, with each replicate consisting of three to four technical replicates. The error bars represent the standard error of the mean. Two-tailed paired t-tests were performed to compare the mCherry signal in the knockout cells to the signal in control knockout cells. *p < 0.05, **p < 0.01, ***p < 0.001, and ****p < 0.0001. “ns” indicates not significant.

(B) Cells transfected with the indicated sgRNAs were infected with SINV at an intermediate MOI (10-fold more viral suspension than in the mCherry fluorescence measurement experiments in Panel A). Thirty hours later, the cell culture medium was collected and filtered through a 0.45 µm filter. It was then centrifuged overnight at 3,500 g and 4°C. The viral particles were then resuspended in 100 µL of PBS, and an equal volume of each suspension was used to evaluate the production of progeny virions by immunoblotting.

(C) HEK293T cells were infected with viral suspensions prepared in Panel B. The cells were then collected 24 hours later. The mCherry fluorescence was quantified by flow cytometry. Representative profiles are shown.

(D) Cells that were transfected with the indicated sgRNAs were infected with SINV at a high MOI for 12 hours. An immunoblot was performed using an antibody against the spliced XBP1 isoform, a marker of ER stress. For positive controls, cells were treated with the ER stress inducers tunicamycin (5 µg/mL) or thapsigargin (1 µM) for five hours, or with the vehicle.

(E) Bar plot showing the quantification of mCherry fluorescence in cells in which the DENR or EIF5A genes were edited using a second set of sgRNAs. The bar plot represents the mean of three biological replicates, with each replicate consisting of three to four technical replicates. The error bars and statistical tests are as described in Panel A.

**Extended data Figure 6.**

(A) Cells transfected with the indicated sgRNAs were infected with SINV at a MOI corresponding to 10-fold and 2-fold more viral suspension than in the mCherry fluorescence measurement experiments shown in Figure 5A. Thirty hours later, the cell culture medium was collected, filtered through a 0.45 µm filter, and centrifuged overnight at 3,500 g and 4°C. The viral particles were then resuspended in 100 µL of PBS, and an equal volume of each suspension was used to evaluate the production of progeny virions by immunoblotting.

(B) Immunoblot analysis of phosphorylated and total eiF2α in cells transfected with control or *PKR*-targeting sgRNAs. Cells were either mock-infected or infected with SINV at a high MOI for 13 hours. An immunoblot against PKR shows the efficiency of the knockout.

**Extended data Figure 7.**

(A) Scatterplot showing the relationship between changes in protein abundance in *ASCC3*-knockout cells that were either mock-infected (x-axis) or infected with SINV at a high MOI for 13 hours (y-axis). Proteins containing a signal peptide are colored turquoise, and ASCC3 is colored fuchsia.

(B) Scatterplot showing a comparison of changes in protein abundance in cells for which two distinct sgRNAs were used for *ASCC3* gene editing. Proteins containing a signal peptide are colored turquoise. ASCC3 and ASCC2 are colored fuchsia.

(C) Scatterplot showing the relationship between changes in protein abundance in *ZNF598*-knockout cells (x-axis) and *ASCC3*-knockout cells using the second sgRNA for *ASCC3* gene editing (y-axis). Proteins containing a signal peptide are colored turquoise. ZNF598, ASCC3, and ASCC2 are colored fuchsia.
